# Benchmarking of a Bayesian single cell RNAseq differential gene expression test for dose-response study designs

**DOI:** 10.1101/2021.09.08.459475

**Authors:** Rance Nault, Satabdi Saha, Sudin Bhattacharya, Jack Dodson, Samiran Sinha, Tapabrata Maiti, Tim Zacharewski

**Author notes:** Co-corresponding authors: Tim Zacharewski, Michigan State University, 1129 Farm Lane, Rm 248. East Lansing, MI 48824, Tapabrata Maiti, Michigan State University, 619 Red Cedar Rd. C415 Wells Hall. East Lansing, MI 48824. Equal contribution (Rance Nault, Satabdi Saha).

## Abstract

The application of single-cell RNA sequencing (scRNAseq) for the evaluation of chemicals, drugs, and food contaminants presents the opportunity to consider cellular heterogeneity in pharmacological and toxicological responses. Current differential gene expression analysis (DGEA) methods focus primarily on two group comparisons, not multi-group dose-response study designs used in safety assessments. To benchmark DGEA methods for dose-response scRNAseq experiments, we proposed a multiplicity corrected Bayesian testing approach and compare it against 8 other methods including two frequentist fit-for-purpose tests using simulated and experimental data. Our Bayesian test method outperformed all other tests for a broad range of accuracy metrics including control of false positive error rates. Most notable, the fit-for-purpose and standard multiple group DGEA methods were superior to the two group scRNAseq methods for dose-response study designs. Collectively, our benchmarking of DGEA methods demonstrates the importance in considering study design when determining the most appropriate test methods.

## Introduction

Single-cell transcriptomics enables researchers to investigate homeostasis, development, and disease at unprecedented cellular resolution^1-5^. As with any new innovative technology, diverse tools soon follow to address specific applications and unique challenges. Currently, there are dozens of differential gene expression analysis (DGEA) approaches for single-cell RNAseq (scRNAseq) data; developed based on differences in assumptions, statistical methodologies, and study designs^6-11^. A recent comparison of 36 approaches demonstrated acceptable performance for common bulk RNAseq tools such as edgeR and limma-trend, and MAST for snRNAseq, as well as common statistical tests such as the Wilcoxon Rank Sum (WRS) and the t-test^9^. However, most methods have been developed primarily for two group comparisons whereas experiments that include multiple groups, such as when assessing risk in pharmacology and toxicology studies where dose-response designs are required. The use of two sample tests for multiple group study designs elevate the type I error rate warranting further investigation of these methods for multiple group dose-response study designs^12^.

Dose-response studies are used to derive the efficacy and/or safety margins such as effective dose and the point of departure (POD). Significant efforts by the toxicology and regulatory communities have suggested that acute (<14 days) and sub-acute (14 – 28 days) transcriptomic studies as viable alternative to the current standard 2-year rodent bioassay that significantly reduces the time and resources needed to assess risk^13-15^. Gene expression profiling at single-cell resolution could further support such evaluations by identifying cell-specific dose-dependent responses indicative of an adverse event. The U.S. National Toxicology Program (NTP) recently reported a robust DGEA approach is essential to deriving biologically relevant PODs^15^. However, concerns regarding the inclusion of false positives that produce less conservative POD estimates potentially leads to incorrect classification of mode-of-action, thus highlighting the importance of controlling type I error rates^16, 17^.

Unlike microarray and bulk RNAseq datasets, single-cell RNAseq (scRNAseq) data is zero inflated due to the low per cell RNA input, biases in capture and amplification, transcriptional bursts, and other technical factors^18^. Consequently, scRNAseq test methods usually consider the gene expression distribution as a mixture of a zero and a positively (non-zero) expressed population^19-21^. For example, the Seurat Bimod approach tests for differential gene expression using a likelihood ratio test designed for the said mixture population. MAST extends the Seurat Bimod test to a two-part generalized linear model structure capable of incorporating covariates^19, 20^. Given the improved performance of MAST^9, 19, 20^, we hypothesized that multiple group tests developed assuming the same distributional framework would be most favorable for dose-response study designs. Furthermore, a Bayesian approach which considers prior knowledge is anticipated to minimize type I error rates^22, 23^.

We propose a novel, multiplicity corrected, Bayesian multiple group test (scBT) designed exclusively for DGEA in zero inflated continuous data populations, characteristic of dose-response scRNAseq data. Two other fit-for-purpose frequentist multiple group tests are also examined: (1) a multiple group extension of the Seurat Bimod test and (2) a simple extension of test (1) to a generalized linear model framework. The proposed methods are benchmarked against commonly used approaches for DGEA on simulated and real experimental dose-response datasets.

## Results

### Dose-response single-cell data simulations

For benchmarking of DGEA methods, a ground truth is required. Existing simulation tools such as PowSimR, SymSim, SPsimSeq, and Splatter are commonly used for power analyses, evaluating DE analysis methods, and testing cell clustering strategies^24-27^. Tools such as SymSim and Splatter are also capable of simulating cell trajectories and model differentiation processes. Trajectories which exhibit non-linear changes over time or across different developmental stages are not unlike dose-response effects which change over a continuum of doses. However, dose-responsive changes commonly follow defined trajectories such as Hill, exponential, power, and linear models^28^. To simulate dose-response scRNAseq data we developed a wrapper for the Splatter scRNAseq data simulation tool named SplattDR. SplattDR modified the Splatter grouped data simulation strategy by adjusting counts from means defined by one of the dose-response functions outlined in the Materials and Methods.

To demonstrate the modeling capability of SplattDR, 10,000 gene expression responses were simulated with a 10% probability of being differentially expressed, equally distributed across the dose-response models. Parameters used in Splatter were initially estimated from our experimental single nuclei RNAseq (snRNAseq) dose-response dataset. The simulated data compared to the experimental data showed the relationship between the mean expression, percentage of zeroes, and mean variance were consistent (**Figs. 1a-b**). Estimation of the normalized root mean square deviation (NRMSD) from a curve fit to the experimental data indicated excellent concordance. This strong concordance was also maintained within distinct dose groups (**Figs. S1-2**). The distribution of log(fold-changes) between vehicle (dose 0) and the highest simulated dose (dose 9; 30 µg/kg) showed a more even distribution within a similar range compared to experimental data which was skewed towards induction (**Fig. 1c**). However, the gene induction skew was captured by modulating the parameters affecting the probability of differential expression and the proportion of differentially repressed genes (**Fig. S3**). Principal components analysis (PCA) of the simulated data clearly showed the dose-dependent characteristics of scRNAseq data with distinct clusters increasing in separation with increasing dose (**Fig. 1d**) which was also resolved by PCA within the experimental data (**Fig. S4)**.

**Fig. 1:**
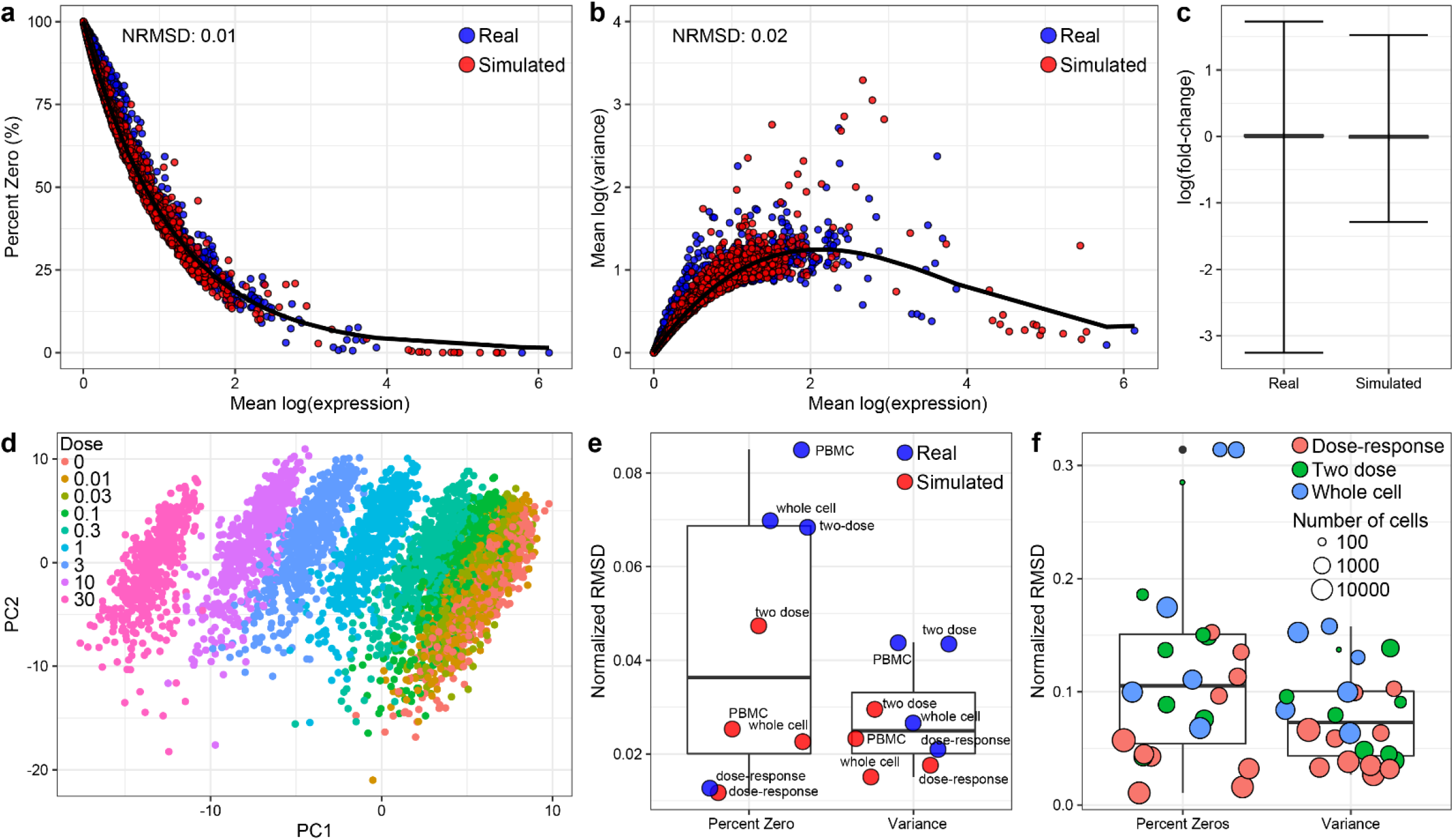
Comparison of simulated and real dose-response data. **a** Relationship between gene-wise mean expression and percent zeroes for simulated and real dose-response data. Simulation data consisted of 10,000 genes and 9 dose groups based on parameters derived from experimental dose-response snRNAseq data. Black line represents a fitted model to the experimental data from which the normalized root mean square deviation (NRMSD) of simulated data was determined. **b** Relationship between gene-wise mean expression and variance for simulated and experimental data. NMRSD was calculated for simulated data from the fitted model represented as a black line. **c** Distribution of log(fold-changes) in experimental and simulated data showing the median and minimum and maximum values. **d** Principal components analysis of simulated data colored according to simulated dose groups. **e** NMRSD estimated relative to fitted model in **a**,**b** for simulated data generated from initial parameters derived from published hepatic scRNAseq (two dose; GSE148339), hepatic whole cell (whole cell; GSE129516), and peripheral blood mononuclear cell (PBMC; GSE10’8313) datasets. **f** NMRSD estimated relative to model fitted to cell-type specific experimental dose-response data when simulated from initial parameters estimated from that same cell type. Box and whisker plots show median NMRSD, 25 and 75^th^ percentiles, and minimum and maximum values.

To our knowledge, no other published *in-vivo* dose-response scRNAseq datasets are available limiting the number of datasets to estimate initial parameters for simulation to date. To investigate whether existing datasets generated using a different study design (*e*.*g*., whole cells or different tissue source) could be used to derive initial parameters, we also simulated 10,000 genes starting with parameters estimated from (i) a two-dose liver snRNAseq (GSE148339), (ii) whole cell liver scRNAseq (GSE129516), and (iii) peripheral blood mononuclear cells (PBMC; GSE108313) datasets. When compared to a model fit for experimental data to determine the relation between mean expression and percent zeroes or mean variance, the NRMSD for data simulated from these datasets were between 1 - 10% with data simulated from whole cell data differing the most from the model fit (**Fig. 1e**). We then explored whether parameters estimated from distinct cell types could replicate the characteristics of that same cell type (**Fig. 1f**). Not surprisingly, lower NRMSD values were observed for simulated cell-specific data based on experimental dose-response data estimated starting parameters with whole cell data performing the worst. Notably, when data derived from a lower abundant cell subtype was used to estimate starting parameters, the dose-response characteristics for that cell subtype was also poorly modeled (**Figs. 1e-f, S1-2**).

### Performance Accuracy of DE test methods

We evaluated the performance of several differential gene expression analysis methods on simulated datasets consisting of 9 dose groups of 500 cells each (4,500 total) and 5,000 genes with a 10% probability of being differentially expressed (500 differentially expressed genes). Selection criteria for test inclusion are outlined in the Materials and Methods section and included 9 test methods; ANOVA^29^, single-cell Bayes Hurdle Model test (scBT), Kruskall-Wallis (KW)^30^, limma-trend^31, 32^, Likelihood-ratio test (LRT) linear and multiple, MAST^19^, Seurat Bimod^33^, and Wilcoxon Rank Sum (WRS)^34^. With ground truth from simulated data, the sensitivity, specificity, and precision for each test method was computed. Area under the receiver-operating characteristic curve (AUROC) was used to measure test performance for correctly classified differentially expressed genes. In unfiltered data, AUROC scores showed similar performance for most tests except scBT which had the largest AUROC among all test methods (**Fig. 2a**). To account for the inherent class imbalance between differentially expressed and non-differentially expressed classes the area under the precision-recall curves (AUPRC) was also calculated. Similar to AUROCs, AUPRCs identified scBT as the best performing test (**Fig. 2c**). In most standard differential expression testing pipelines genes expressed at low levels are removed to minimize false detection rates. Following filtering of genes expressed in ≤5% of cells in any dose group, scBT was consistently ranked as the best test based on AUROC and AUPRC scores. The performance of LRT linear test also improved, with comparable AUROC and AUPRC scores relative to scBT, suggesting LRT linear is poorly suited for genes expressed at low levels (**Fig. 2b,d**).

**Fig. 2:**
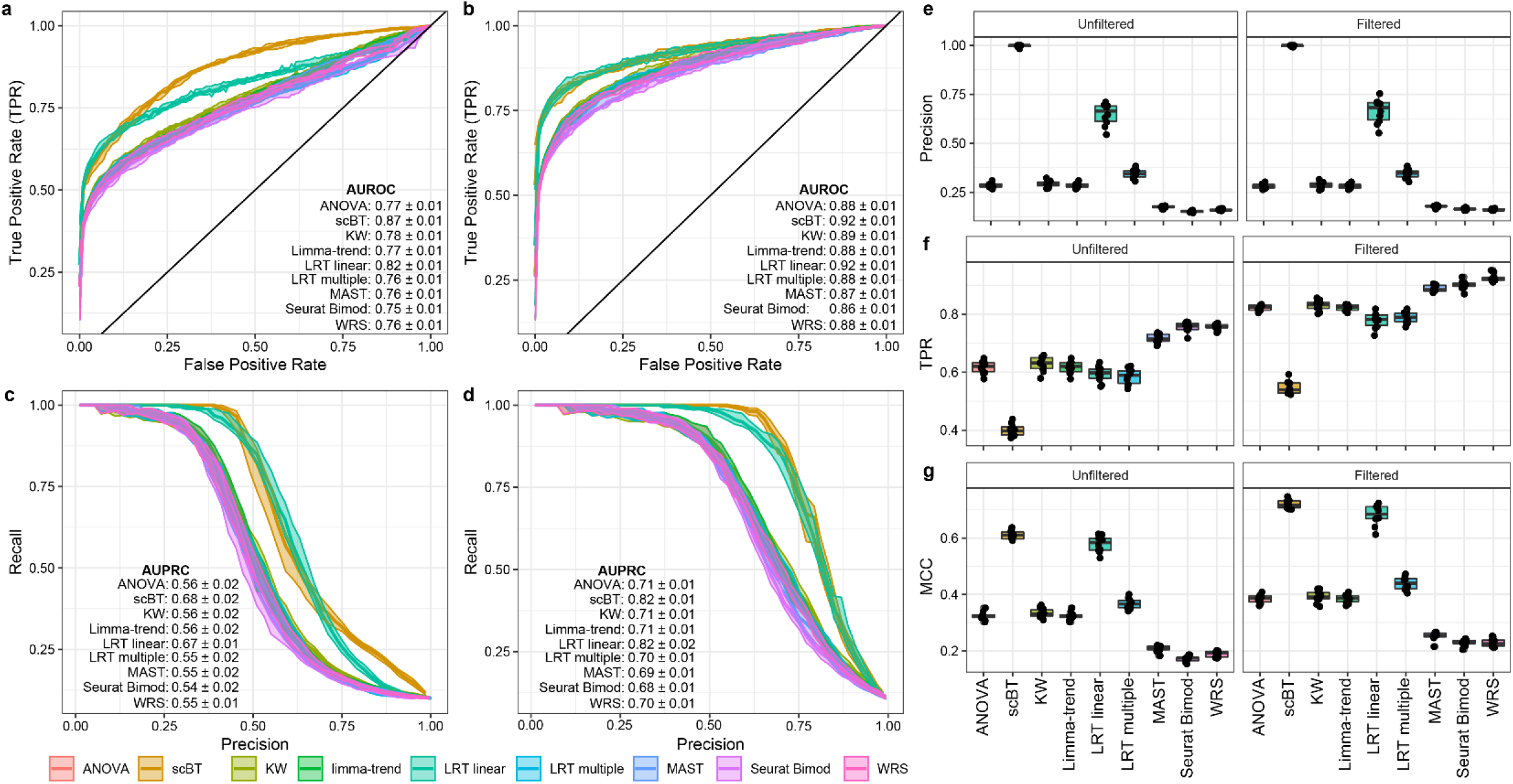
Classification performed of DE analysis tests. **a** ROCs estimated from simulated dose-response scRNAseq data for 9 DE test methods including all genes expressed in at least 1 cell (unfiltered). **b** ROCs for 9 DE test methods after filtering simulated dose-response scRNAseq data for genes expressed in only ≥5% of cells (low levels) in at least one dose group. **c** Precision-recall curves (PRCs) for 9 DE test methods on unfiltered simulated dose-response scRNAseq data. **d** PRCs for 9 DE test methods on filtered simulated dose-response scRNAseq data. Lines represent the mean values and shaded region reflects the standard deviation for 10 independent simulations. **e** Precision of DE test methods. **f** FPR of DE test methods. **g** MCC for test methods. **e,f,g** Box and whisker plots median values, 25^th^ and 75^th^ percentiles, and minimum and maximum values for 10 independent simulations. Points reflects values for each independent simulation. Panels display comparisons of unfiltered and filtered datasets.

AUROC and AUPRC reflect the performance of each test method with varying significance (*i*.*e*., *p*-value) thresholds. In the standard pipeline a fixed threshold is used, typically a *p*-value ≤ 0.05 after adjustment for multiple hypothesis testing (*i*.*e*., Bonferroni correction). For each method except scBT, the performance at an adjusted *p*-value ≤ 0.05 significance criteria was evaluated. In scBT analysis, a gene was considered differentially expressed when the estimated posterior probabilities of the null hypothesis, *p*(*H*_0,*j*_|*D*_*j*_), was less than *ζ*, where the *ζ* value was chosen to achieve a target FDR of 0.05. scBT significantly outperformed all other tests in precision rates irrespective of low expression filtering (**Figs. 2e, S5**). However, scBT was less effective in identifying true positives (**Figs. 2f, S5**). Applying the filtering criteria improved the recall rates, but the precision rates remain largely unchanged (**Figs. 2e,f**). Test method classification performance scores were estimated as the Matthews Correlation Coefficient (MCC) which is well suited for unbalanced data^35^. We see that the scBT and LRT linear tests performed best for this metric on both unfiltered and filtered data (**Fig. 2g**).

### Type I error control and power

To investigate test performance in controlling type I errors (false positives), DGEA methods on simulated datasets were examined with 0% DE genes (*i*.*e*., negative control). Using the ζ threshold for the computed posterior null probabilities, scBT identified only 1 false positive gene in 2 of 10 simulations (**Fig. 3a**). ANOVA, scBT, KW, limma-trend, and LRT linear had false positive rates (FPRs) below 3% indicating better performance compared to two group tests. After filtering for genes with low expression levels, scBT still correctly identified all the non-differentially expressed genes and was the best performing test. These are the same tests that had a better FPR control in initial simulations (**Fig. 2**). To explore whether mean expression or percentage of zeroes influenced type I error rates, a logistic regression model was fit to negative control data. We predicted the probability for each gene to be identified as differentially expressed in the negative control data. While the curve for scBT is missing since few false positives were identified, the predicted FPR for all the other tests except LRT linear were also high for highly expressed genes with few zeroes (**Fig. 3b,c**). Next, a positive control dataset with 100% differentially expressed genes was simulated to evaluate test performance for detecting true positives. All tests except scBT exhibited a false negative rate (FNR) ≤ 40% (**Fig. 3d**). The best performing tests for FNR also had high FPR. Logistic model regression fitting for false negative classification of genes shows that the false negative rates were highest when the mean expression was either too high or too low for all tests (**Fig. 3e,f**).

**Fig. 3:**
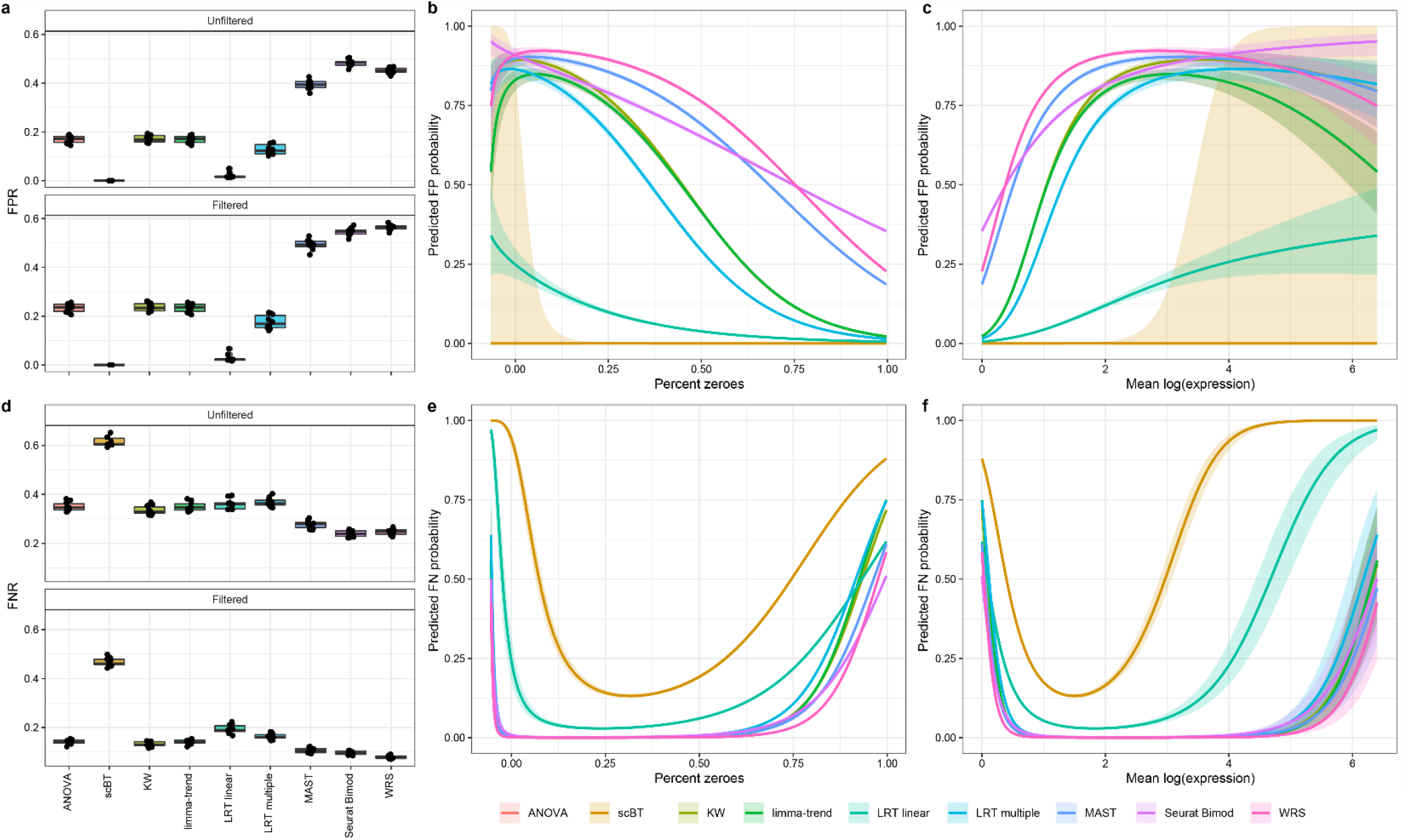
Evaluation of Type I and II error control. **a** False positive rate (FPR) of 9 differential expression test methods estimated from negative control (0% DE genes) simulated dose-response scRNAseq data including all genes expressed in at least 1 cell (unfiltered) and genes expressed in only ≥ 5% of cells in at least one dose group (filtered). **b,c** Logistic regression models were fitted to negative control data to predict the probability of false positive identification using percent zeroes and mean expression as covariates. Lines represent the predicted probability of false positive classification with the shaded region representing the 95% confidence interval. **d** False negative rate (FNR) of 9 differential expression test methods estimated from positive control (100% DE genes) simulated dose-response scRNAseq data including unfiltered and filtered datasets. **e,f** Logistic regression models were fit to positive control data. Lines represent predicted probability of false negative classification with shaded region representing the 95% confidence interval.

### Parameter sensitivity analyses

Experimental scRNAseq datasets will vary between cell types, cell composition, and responses depending on the target tissue, treatment, number of cells sequenced, and more. For example, some distinct cell types are very abundant (*e*.*g*., hepatocytes), with others present at lower levels (*e*.*g*., portal fibroblasts) in hepatic scRNAseq datasets. Moreover, treatments such as exposure to a xenobiotic, can elicit dose-dependent changes in relative proportions of cell types such as the infiltration of immune cells^36^. We investigated the impact by changing cell abundance from 25 to 2,000 cells per dose group and observed an increase in the false positive rate (FPR) when increasing the number of cells (**Fig. S6**). The scBT and LRT linear tests were less sensitive to an increase in the FPR as cell abundance increased while the total positive rates (TPR + FPR) increased with cell abundance for all methods. Although all tests exhibited comparable performance at low cell numbers (≤ 500), as cell numbers increased scBT outperformed all other tests in both precision and MCC score (**Fig. 4a, S6**). Comparison of AUROCs and AUPRCs across cell numbers showed that ANOVA, KW, limma-trend, and LRT linear tests performed best for a small number of cells, but the increase in AUROC was steeper for scBT (**Fig. S7-8**).

**Fig. 4:**
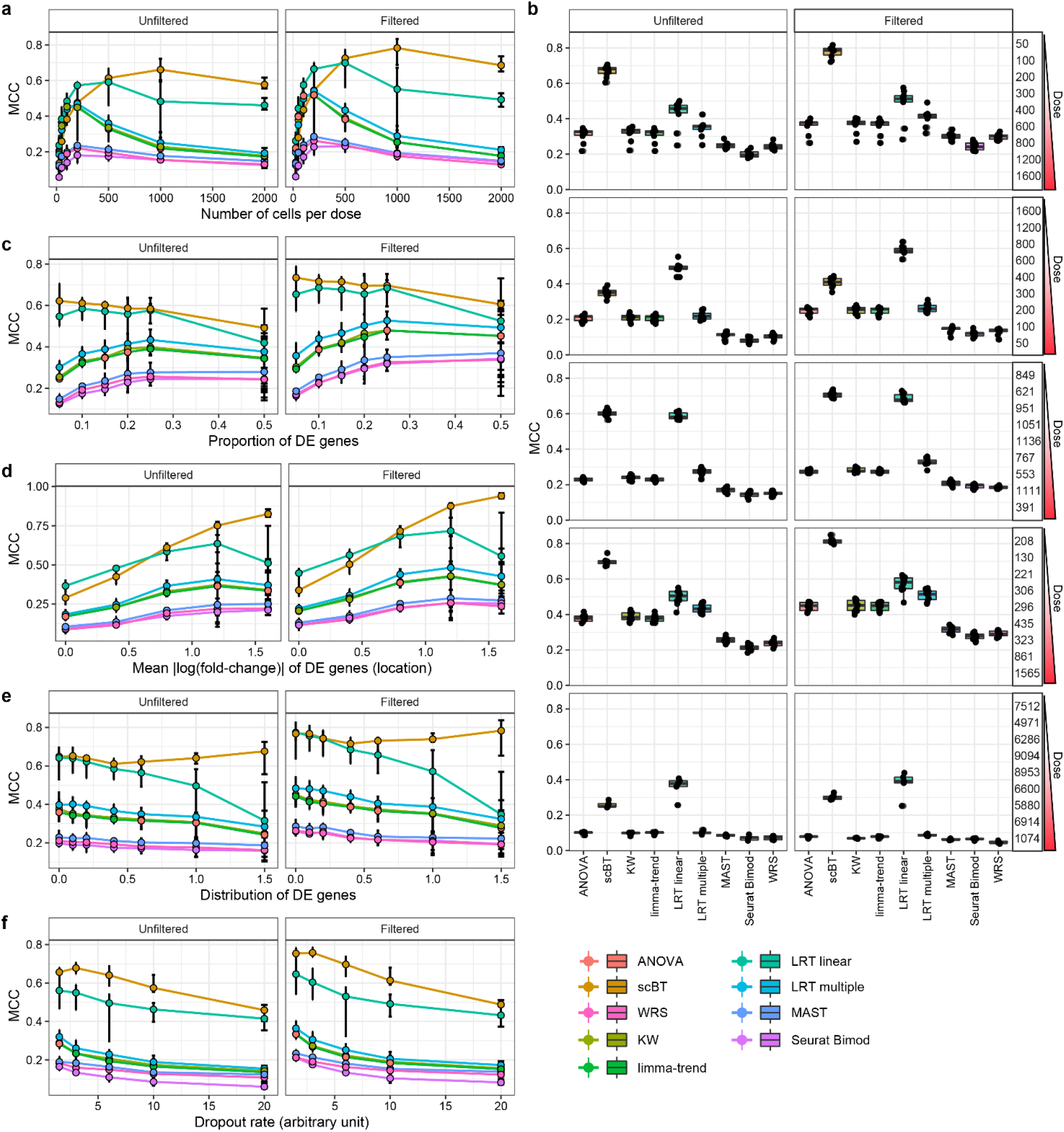
Matthews correlation coefficient (MCC) from sensitivity analyses of differential expression test methods. **a** MCC for 9 DGEA test methods determined from simulated dose response data with varying number of cells per dose group. Simulations consisted of 5,000 genes with a probability of differential expression of 10% and 9 dose groups. **b** MCC for simulated data varying the cells numbers by dose group. The number of cells in each of the 9 doses groups is shown on the right. **c** MCC for varying proportion of differentially expressed genes. **d** MCC for varying proportion of repressed differentially expressed genes. **e** MCC for varying fold-change location. **f** MCC for varying fold-change scale. Points represent median and error bars represent minimum to maximum values. Boxplots represent median, 25^th^ to 75^th^ percentile, and minimum to maximum values. Each analysis consisted of 10 replicate datasets including all genes expressed in at least 1 cell (unfiltered) and genes expressed in ≥ 5% of cells in at least one dose group (filtered).

It was also evident from the experimental snRNAseq dataset that the number of cells per dose group was not fixed. We evaluated the performance of the test methods when the number of cells dose-dependently increased or decreased, and when the number of cells per dose group were taken from experimental data. Notably, while scBT had the best MCC for increasing number of cells per dose, LRT linear performed better than scBT when the number of cells decreased before and after filtering for genes expressed at low levels (**Fig. 4b**). The shift in MCC between increasing and decreasing cell numbers for scBT appears to be driven by a concomitant decrease in FTPR and increase in FNR (**Fig. S9**).

Unique chemical, drug, environmental contaminant, and natural product classes elicit distinct differential gene expression profiles defined by the mode-of-action (MoA) as well as by their metabolism, potency (sensitivity) and efficacy (maximal response). Differences between compound classes are reflected in the gene expression profile in (*a*) the proportion of differential expressed genes, (*b*) the number of induced/repressed genes, (*c*) the mean fold-change for differentially expressed genes, and (*d*) the distribution of fold-change for differentially expressed genes. These 4 parameters were modulated in simulated data to determine the effect of the percentage of differentially expressed genes, the mean fold-change (aka *location*), and the fold-change distribution (aka *scale*) on test performance. Among these scenarios, changing the proportion of repressed genes had little to no impact on test method performance (**Fig. 4c-f, S14**).

Increasing the proportion of differentially expressed genes led to an improvement in MCC except for scBT and LRT linear, though these tests maintained the top MCC scores as well as AUROC and AUPRC (**Fig. 4c, S11-13**). As the magnitude of the effect increased, LRT linear performed best at the low end while scBT exhibited the greatest improvement in MCC (**Fig. 4d**). Conversely, while the MCC decreased for most tests when modulating the fold-change scale of differentially expressed genes, scBT improved and was more stable (**Fig. 4e**). As zero inflation increased, the FPR increased and the precision decreased for all tests (**Fig. S23**). However, scBT was least affected, and maintained the highest MCC among all tests (**Fig. 4f**). AUROC and AUPRC values also indicated that scBT consistently outperformed other test methods (**Fig. S24-25**).

### Test method agreement

To assess agreement between tests, the area under the concordance curve (AUCC) for each pair of tests for the top 100 genes ranked by adjusted *p*-value was calculated as previously described^9, 37^. All methods showed excellent concordance (AUCC ≥ 0.77) with LRT linear showing the poorest consistency compared to all other tests while the limma-trend and ANOVA tests showed perfect agreement with an AUCC of 1 (**Fig. S5**). Pairwise differential gene expression comparisons between DE, Seurat Bimod, MAST and WRS had AUCCs >0.95 AUCCs while the multiple group tests ANOVA, LRT multiple, KW, and scBT clustered together with AUCC ranging between 0.9 - 1. In the absence of nuisance covariates, MAST and Seurat Bimod provided similar results, as expected given their similar mixture normal model structure. Likewise for ANOVA and limma-trend, both of which rely on normality assumptions for testing differential gene expression.

### Real dose-response dataset DE analysis

Without ground truth for experimental data, the performance of the differential expression test methods was examined by first evaluating the agreement for each identified cell type (**Fig. 5, S26**). Genes in the experimental dataset were considered differentially expressed when expressed in ≥5% of cells in at least one dose group and had a |fold-change| ≥ 1.5. In hepatocytes, the most abundant cell type, fewer than 5 genes were not detected in all test methods, with the majority missed by the WRS test (**Fig. 5a**). Upon closer examination, those genes were not expressed in control hepatocytes. Not surprisingly, for all cell types, the largest intersection was between all tests indicating strong agreement within all test methods. Only a few tests identified a subset of unique genes as differentially expressed, which accounted for a very small fraction. For example, LRT linear identified 12 unique differentially expressed genes in portal fibroblasts one of the least abundant cell types (**Fig. 5b**). LRT linear was the best performing test for low cell numbers indicating that the 12 unique differentially expressed genes may in fact be true positives. Consistent with simulations of varying cell numbers (**Fig. 4a**), 24 genes were not identified as differentially expressed by the scBT method for stellate cells which exhibit a dose-dependent decrease in numbers (**Fig. 5c,d**). Although scBT outperformed other tests in most scenarios, it under performed in this scenario. Nevertheless, when ranking genes by significance level (*i*.*e*., *p*-values), AUCC were high for all pairwise comparisons.

**Fig. 5:**
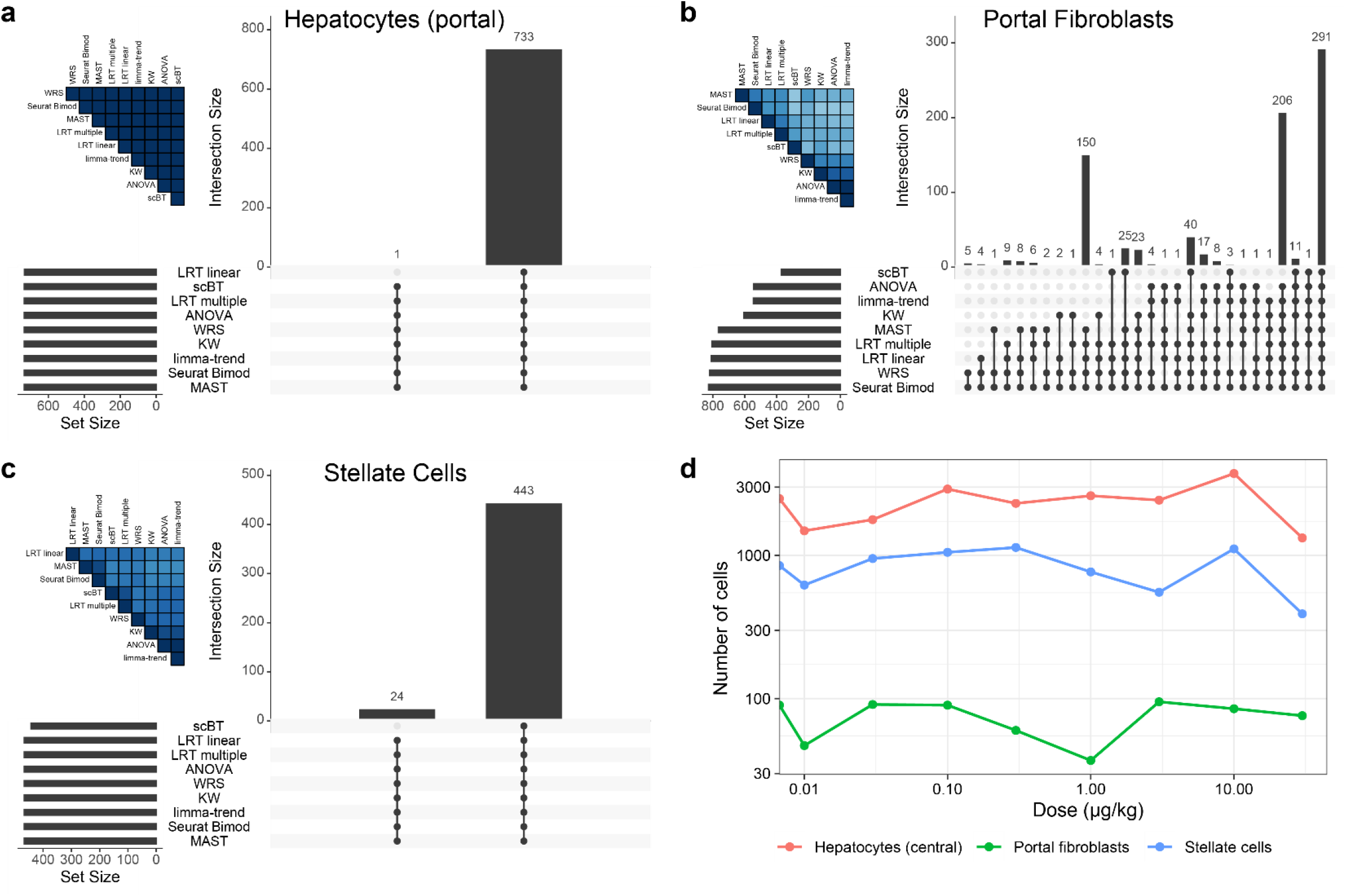
Agreement of differential expression test methods on experimental dose-response data. **a** Upset plot showing the intersection size of genes identified as differentially expressed by 9 different test methods in hepatocytes from the portal region of the liver lobule. **b** Intersect of differentially expressed genes in portal fibroblasts. **c** Intersect size in hepatic stellate cells. Vertical bars represent the intersect size for test methods denoted by a black dot. Horizontal bars show the total number of differentially expressed genes identified within each test (set sizes). Only intersects for which genes were identified are shown. Genes were considered differentially expressed when (i) expressed in >5% of cells within any given dose group and (ii) exhibit a |fold-change| ≥ 1.5. A heatmap in the upper left corner of each panel shows the pairwise AUCC comparisons for the 500 lowest *p*-values. **d** Relative proportion of cell types identified in each dose group of the real dataset for the cell types in **a,b,c**. Experimental snRNAseq data was obtained from male mice gavaged with sesame oil vehicle (vehicle control) or 0.01 – 30 µg/kg TCDD every 4 days for 28 days.

## Discussion

The goal of this study was to compare the performance of newly developed DGEA test methods for dose-response experiments to existing analysis methods. Using simulated data to generate ground truth, we evaluated the performance of 9 differential expression testing methods which were broadly classified as either fit-for-purpose, multiple group, or two group tests. Criteria for test method selection was based on previous benchmarking efforts for two group study designs identifying MAST, limma-trend, WRS, and t-test as the best performers^9, 38^. ANOVA and KW tests were also included for evaluating multiple group comparisons, and Seurat Bimod, for having the same modelling framework as scBT, LRT multiple, and LRT linear tests. The test methods were ranked from best to worse (1 – 9) based on type I error rate, type II error rate, MCC, AUROC, and AUPRC (**Fig. 6, Table S1**).

**Fig. 6:**
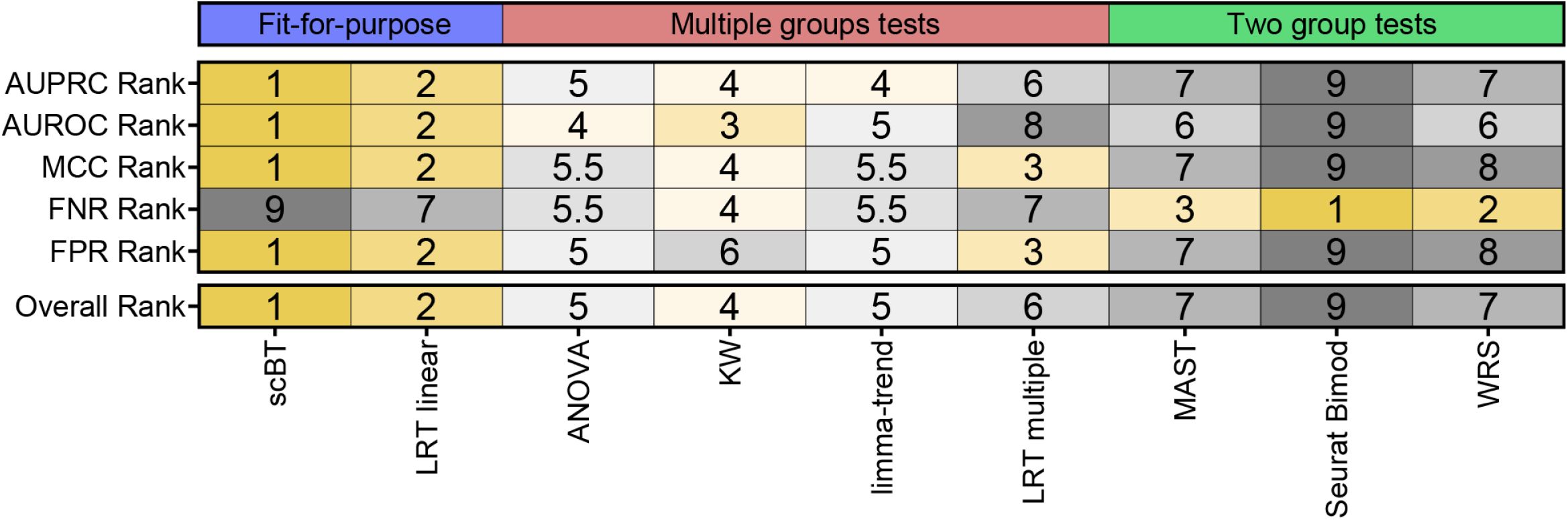
Median ranking of differential expression test methods across all simulations. **The** median rank of each test method was calculated for AUPRC, AUROC, MCC, FNR, and FPR. Tests were grouped according to intended application including fit-for-purpose tests developed for the analysis of dose-response datasets, multiple group tests, and two group tests. The overall rank represents the median value for the 5 key metrics presented here.

While several scRNAseq tools have been developed^24-27^, none are developed to simulate dose-response models commonly identified in toxicological and pharmacological datasets^28, 39^. Our SplattDR wrapper for the Splatter package^27^ was able to show that simulated data can effectively emulate key experimental scRNAseq data characteristics when simulation parameters were estimated from various Unique Molecular Identifier (UMI)-based datasets. In agreement with a previous report, technical and biological factors, such as cell type, does appear to influence gene dropout rates^18^. We primarily focused on 10X Genomics UMI data given the unavailability of real experimental dose-response data generated using other platforms.

Overall, test method performance was consistent with their intended application. For example, fit-for-purpose tests scBT and LRT linear consistently ranked higher followed by multiple groups tests such as KW and LRT multiple. scBT exhibited the best overall performance with excellent FPR control and top ranked MCC while LRT linear struck a balance between type I and type II error rates. The scBT results are not surprising as Bayes factor-based tests have proven to be conservative and consequently more appropriate when false positives are of concern^22, 23^. In the context of investigating chemical or drug MoAs, false positives have the potential to lead to wasted effort and resources in attempts to validation and support findings^40^. Moreover, when assessing a large number of genes, a 5% FP rate (*P*-value ≤ 0.05) can result in hundreds of FPs that skew MoA classifications^17^.

A single test method was not expected to outperform all other tests under all conditions as previously demonstrated when comparing pairwise testing^6, 9, 38^. Therefore, we assessed the strengths and limitations of each test method by varying parameters likely to change within and across various experimental datasets. The number and relative abundance of cell types is known to be affected by disease or treatment, and the distribution of differential expression influenced by the chemical, drug, or food contaminant being evaluated^5, 36^. scBT consistently ranked at the top under most scenarios, particularly when the mean and standard deviation of the fold-change for differentially expressed genes varied. However, scBT under performed in MCC when the number of cells decrease in a dose-dependent manner which would be expected in treatments which alter cell population sizes (*e*.*g*., inflammation). Under these circumstances LRT linear outperformed all other tests with scBT performing similar to the other test methods as evident when 24 differentially expressed genes were not identified by scBT within experimental data for stellate cells which experienced a dose-dependent decrease in relative abundance following TCDD treatment. Although excluding genes expressed at low levels generally improved the performance of all test methods, the comparative performance of test methods did not significantly change in most cases.

Collectively, our findings suggest that scBT and LRT linear fit-for-purpose tests are better suited for the differential expression analysis of dose-response studies and when false positives are of greater concern than false negatives. Moreover, consistent with previous benchmarking efforts, we show that common non-parametric tests such as KW outperform test methods developed for scRNAseq data when the study involves comparisons between multiple groups. Ultimately, each test method performs optimally under diverse scenarios. While the importance of controlling type I error rates is acknowledged, a balance must be struck with type II error rates. The tradeoff should be determined based on the individual research question being investigated. It may even become reasonable to apply disparate test methods for distinct cell types based on dropout rates, cell abundance, and changes in relative cell proportions given the strengths and weaknesses of each test method.

## Methods

### Animal handling and treatment

Animal handling and treatment Male C57BL/6 mice aged postnatal day (PND) 25 were obtained from Charles Rivers Laboratories (Kingston, NY) were housed and treated as previously described^41^. Mice were housed in lnnovive cages (San Diego, CA) with ALPHA-dri bedding (Shepherd Specialty Papers, IL) at 23°C, 30-40% relative humidity, and a 12:12h ligh:dark cycle. Aquavive water (lnnovive) and Harlan Teklad 22/5 Rodent Diet 8940 (Madison, WI) was provided ad libitum. On PND 29, randomly assigned mice were gavaged at Zeitgeber time (ZT) 0 with 0.1 ml sesame oil vehicle (Sigma-Aldrich, St. Louis, MO), 0.01, 0.03, 0.1, 0.3, 1, 3, 10, or 30 g/kg TCDD every 4 days for 28 days (7 total administered doses). At day 28 mice were euthanized by CO2 asphyxiation and livers were immediately flash frozen in liquid nitrogen and stored at − 80°*C*. All animal procedures were approved by the Michigan State University (MSU) lntitutional Animal Care and Use Committee (IACUC) and reporting of in vivo experiments follow the Animal Research: Reporting of In Vivo Experiments (ARRIVE) and Minimum Information about Animal Toxicology Experiments (MIATE) guidelines (https://fairsharing.org/FAIRsharing.wYScsE).

### Real scRNAseq datasets

Hepatic single-nuclei RNA-sequencing (snRNAseq) was performed as previously described using the 10X Genomics Chromium Single Cell 3’ v3.1 kit (10X Genomics, Pleasanton, CA)^36^. Briefly, nuclei were isolated using EZ Lysis Buffer (Sigma-Aldrich), homogenized by disposable Dounce homogenizer, washed, filtered using a 70-µm cell strainer. The nuclei pellet was resuspended in buffer containing DAPI (10 µg/ml), filtered using a 40-µm strainer, and immediately sorted using a BD FACSAria llu (BD Biosciences, San Jose, CA) at the MSU Pharmacology and Toxicology Flow Cytometry Core (facs.iq.msu.edu/). Sequencing (150-bp paired end) was performed at a depth of 50,000 reads/cell using a NovaSeq6000 at Novogene (Beijing, China). Cell Ranger v3.0.2 was used to align reads to mouse gene models (mm10, release 93) including intrans and exons to consider pre-mRNA and mature mRNA. Seurat was used to integrate and log-normalize expression data^42^. Additional real datasets were publicly available. Hepatic whole-cell generated using the 10X Genomics platform was obtained from GEO (GSE129516) ^5^. Hepatic single-nuclei processed as the dose-response data for control and high dose TCDD treatment (0 and 30 µg/kg) was obtained from GEO (GSE148339). Peripheral blood mononuclear cell (PBMC) data also generated using the 1OX Genomics platform and Seurat was obtained from the SeuratData R package^43^.

### Dose-response data simulation

To simulate dose-response scRNAseq datasets we developed a wrapper for the Splatter R package^27^. Splatter simulates counts using parameters estimated from real data to set the mean expressions, variance, and outlier probability. Other parameters such as the number of cells, genes, probability of being differentially expressed, mean fold-change of DE genes (location) and standard deviation of fold-change of DE genes (scale) were manually assigned to best reflect real data. The wrapper (SplattDR) leverages the group simulation feature of Splatter by applying a multiplicative factor estimated using dose-response models in Table 1 based on the US EPA Benchmark Dose Software^28^. SplattDR R package is available at (github.com/zacharewskilab/splattdr).

**Table 1.**
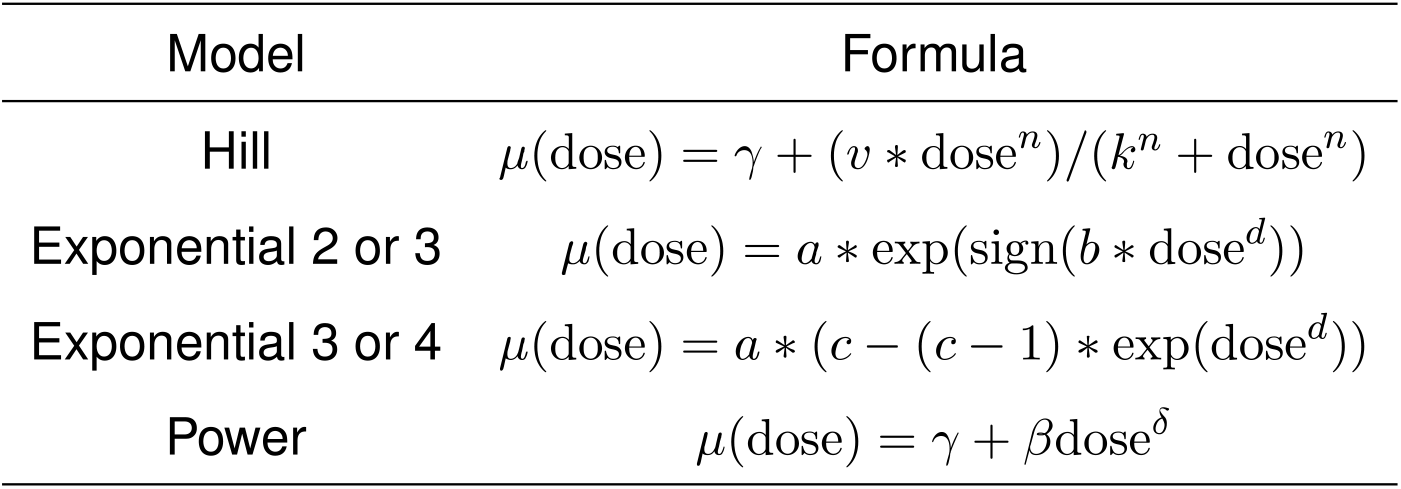
Dose-response models for simulation of scRNAseq data.

### Single cell RNA-seq Hurdle Model

We model the log-normalized gene expression matrix using a hurdle distribution wherein the rate of gene expression is assumed to follow a Bernoulli distribution and conditional on a cell expressing the gene, the log-normalized expression level is assumed to follow a Gaussian distribution^20^ We denote *Y*_*i,j*_ to be the log-normalized expression value of gene *j* in cell *i*, for *i* = 1, … *n* and *j* = 1, … *p*. To characterize the bimodal properties of single cell data, for a given cell, a gene is defined to be either positively expressed or undetected. Define *R*_*ij*_ = *I* [*Y*_*ij*_ > 0] to be the indicator variable denoting the presence or absence of expression for gene *j* in cell *i*. Following^20^ the log-normalized gene expression values are modeled as follows:

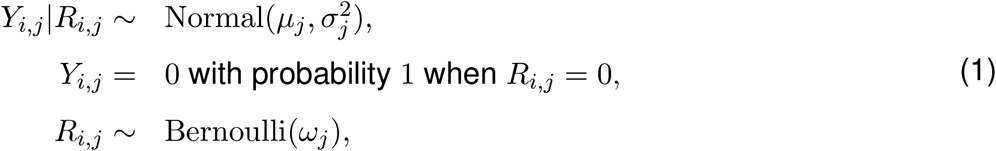

where *µ*_*j*_ and 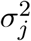 denote the mean and variance of the gene expression level, conditional on the gene being expressed and *ω*_*j*_ denotes the rate of gene expression of gene *j* across all cells.

### Hypothesis Formulation

We now assume that our data has been collected under *K* conditions (doses), and denote the data by*D*_*k,o*_ ≡ {(*Y*_*k,i,j*_,*R*_*k,i,j*_),*i* = 1, …*n*_*k*_}. The underlying populations for the sample data *D*_*k,o*_ for *k* = 1, 2, …, *K*, dose groups are assumed to be identified by the parameters 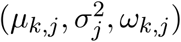. The aim of this study is to test for difference in gene expression patterns between the different dose groups. Traditionally one would perform an ANOVA test to detect changes in mean across groups for samples with continuous measurements. However, to account for the bimodality in single cell gene expression distribution, the test should detect for changes in *µ*_*j*_ and *ω*_*j*_ simultaneously, as both could drive differential gene expression. Therefore we define,

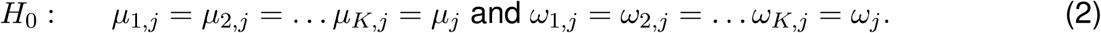

versus the alternative

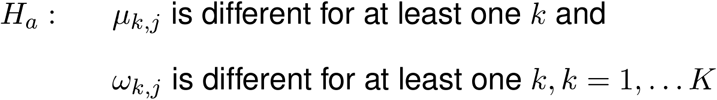

### Single cell Bayesian Hurdle model Analysis (scBT)

Given the single cell RNA-seq hurdle model structure, we sssume that a priori, given 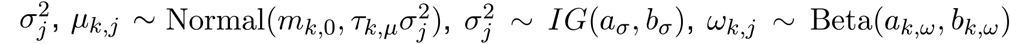, where *IG* is the inverse gamma distribution with shape *a*_*σ*_ and scale *b*_*σ*_ and *m*_*k*,0_, *τ*_*k,μ*_, *a*_*σ*_, *b*_*σ*_, *a*_*k,ω*_, *b*_*k,ω*_ are the hyperparameters. Given the large number of gene-wise model fits arising from a single cell expreriment, there is a pressing need to allow for a parallel structure whereby the same model is fitted to each gene. The prior distributions on the parameters describe how the unknown coefficients *μ*_*k,j*_ *ω*_*k,j*_ and 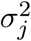 vary across the genes and the dose groups while allowing for information borrowing between the genes. Now, based on the model assumptions, we propose a Bayesian test for simultaneously testing the differences in mean gene expression and dropout proportions as formulated in. Under the null hypothesis the marginal likelihood is written as

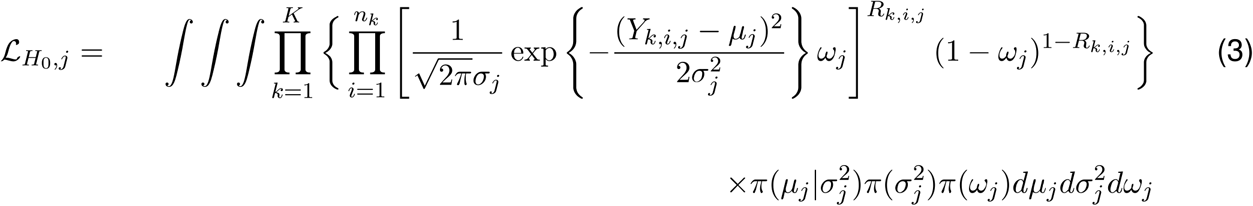

Under the alternative hypothesis we compute the marginal likelihood without any restriction on the *K* means *μ*_*k,j*_ and the dropout parameter *ω*_*k,j*_; *k* = 1, 2, … *K*. Particularly, we assume that 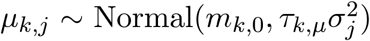, and 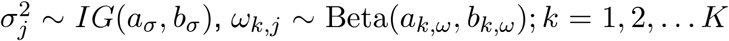. Now, the marginal likelihood under the alternative hypothesis is given by

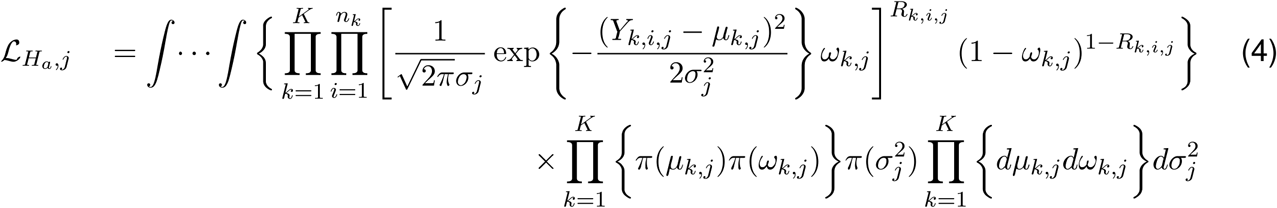

The Bayes factor is then defined as

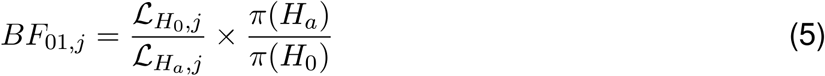

where *π* (*H*_*a*_) and *π* (*H*_0_) are the prior probabilities for the alternative and null model, respectively. The hyperparameters are obtained by maximising the marginal likelihood under the null and the alternative hypothesis. Detailed derivations of the likelihood function and the Bayes Factor are provided in Supplementary Material. Using the test of hypothesis described in Equation (2) scBHM conducts a test of DE for each gene independently. To control for multiplicity we adopt the FDR correction approach discussed in^44^. The rejection threshold is estimated in terms of the posterior probabilities of the null hypothesis, *p*(*H*_0,*j*_|*D*_*j*_). For a target FDR *α*, the procedure rejects all hypotheses with *p*(*H*_0,*j*_ | *D*_*j*_) < *ζ*, where 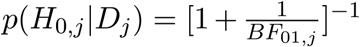 and *ζ* is the largest value such that 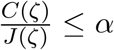 where, *J*(*ζ*) = {*j* : *p*(*H*_0,*j*_|*D*_*j*_) ≤ *ζ*} and *C*(*ζ*) =∑_*j*∈*J*(*ζ*)_ *p*(*H*_0,*j*_|*D*_*j*_).

### Multiple group Likelihood Ratio Test (LRT)

To carry out a direct performance comparison with scBT, we extend the Seurat Bimod^20^ for multiple groups. Assuming that all the K groups have the same variance 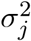 and omitting the index *j* for clarity, the likelihood ratio test can be defined as;

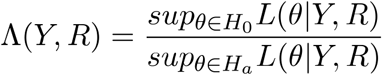

where the likelihood can be written as;

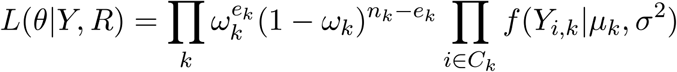

*Y* and *R* represent the gene observation vector and the gene indicator vector across K dose groups, *θ* = {*μ*_*k*_, *σ*^2^, *π*_*k*_, *k* = 1, …, *K*} is the vector of unknown parameters, *C*_*k*_ is the set of cells expressing the gene in group *k* (*i*.*e*.*C*_*k*_ = {*i* : *R*_*i,k*_ = 1}), *e*_*k*_ =∑_*i*_ *R*_*i,k*_ is the number of cells expressing the gene in group *k* and *f* is the density function of the normal distribution with parameters *μ*_*k*_ and *σ*^2^. Following^20^, it can be shown that the combined LRT is the product of a binomial and a normal LRT statistic, both of which can easily be derived using classical statistical theory. Applying classical asymptotic results about LRTs, − 2Iog Λ (*Y, R*) converges to a *χ*^2^ distribution with (2*K* − 2) degrees of freedom under *H*_0_. Detailed derivations of the test statistic can be found in the Supplementary Material.

### Linear model-based Likelihood Ratio Test (LRT linear)

The generalized linear model approach MAST was identified as one of the top performing tests for pairwise differential expression testing^9, 19^. Deriving from their approach, the LRT multiple test is extended to a generalized linear model framework, where the mean and the dropout proportions are modelled as a linear function of the dose groups (assumed to be a continuous covariate). Using the same distributional assumptions defined in (1)we fit a logistic regression model for the discrete variable *R* and a Gaussian linear model for the continuous variable *Y* conditional on (*R* = 1) independently, as follows: *E*(*Y*_*ij*_|*R*_*ij*_ = 1) = *m*_0,*j*_+*m*_1,*j*_ \**d*_*i*_ and *logit*{*P*(*R*_*ij*_ = 1)} = *ψ* _0,*j*_+*ψ*_1,*j*_ * *d*_*i*_, where *d* represents the continuous dose groups. Under this modelling approach, the null hypothesis described in Equation (2) can be rewritten as *H*_0_ : *μ*_*ij*_ = *m*_0*j*_ and logit{*P*(*R*_*i,j*_ = 1)} = *ψ*_0*j*_. The regression models are fit using the *lm* and *brglm* functions in the *stats* and *brglm* R packages. The likelihood ratio test statistic is computed using the same statistical theory discussed discussed for the LRT multiple test and it asymptotically follows a *χ*^2^ distribution under *H*_0_.

### Benchmarking method selection

Our fit-for-purpose tests were benchmarked to existing differential expression testing methods or their multiple group equivalent based on previously reported performance, ability to consider multiple groups, or whether they served as foundation for the scBT and multiple group LRT (LRT multiple) tests developed here. Seurat Bimod served as foundation for the scBT and LRT multiple tests as previously outlined, and MAST was identified as one of the top performing test for two group comparisons^9^. Similarly, limma-trend performed well for two sample comparisons and can consider multiple groups. The Wilcoxon Rank Sum test was identified as providing excellent balance between its ability to identify DE genes and speed, and is the default test for the Seurat R package for scRNAseq analysis. It was also reported that the t test performed well and therefore we included the ANOVA and Kruskal-Wallis (KW) tests, a parametric and non-parametric alternative of the t test for multiple group comparisons. All tests were run without correction for batch effects or other nuisance covariates. Multiplicity for each test was controlled using FDR correction 31. All tests, including scBT, LRT multiple and LRT Linear are available in our scBT R package (github.com/satabdisaha1288/scBT).

### Seurat Bimod

Seurat Bimod test^20^ is a pairwise differential gene expression testing approach developed assuming the single cell RNA-seq hurdle model framework. The test is formulated as *H*_0_, the mean and the dropout parameters of the gene vector under two dose groups are equal versus *H*_*a*_: the mean and the dropout parameters differ over the two groups. The LRT based test statistic −2IogΛ(*y, r*) converges to a *χ*^2^ distribution with (2*K* − 2) degrees of freedom under *H*_0_. The computations are carried out using the R Package *Seurat*.

### MAST

MAST^19^ proposes a two-part generalized linear model for differential expression analysis of scRNAseq data. The first part models the rate of gene expression using logistic regression 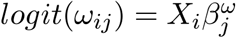 and the second part uses a linear model to express the positive gene-expression *Y*_*ij*_, conditional on *R*_*ij*_ as 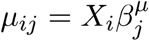; where 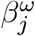 *and* 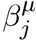 are the coefficients of the covariates used in the logistic and linear regression model respectively. A test with an asympotic *χ*^2^ null distribution is employed for identifying DEGs and multiplicity is controlled using FDR correction^45^. Despite the fact that LAT-linear and MAST have the same hurdle regression framework, the estimation process for the two methods has some significant differences. First, to achieve shrinkage of the continuous variance, MAST assumes a gamma prior distribution on the precision (inverse of variance) parameter and estimates its posterior maximum likelihood estimator (MLE) and uses that in place of the regular MLE of the precision parameter. Second, it fits a Bayesian logistic regression model for the discrete component by assuming Cauchy distribution priors centered at zero for the regression parameters. This is done to deal with cases of “linear separation” where the parameter estimates diverge to ± ∞ and the Fisher information matrix becomes singular. And finally, it considers the cellular detection rate defined as 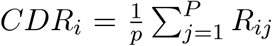 to be a covariate in both the logistic and linear regression models. LRT linear on the other hand simply fits the non-Bayesian linear and the logistic regression models without considering variance shrinkage or adjustment for additional covariates.

### limma-trend

Limma-trend^31^ proposes a linear model based differential expression approach for modelling RNA-seq experiments of arbitrary complexity. Their framework models the mean gene expression as a function of several continuous and categorical covariates. A separate linear model is fitted for each gene, but the gene-wise models are linked by global parameters using the parametric empirical Bayes approaches^32^. The global variance estimated by the empirical Bayes procedure also incorporates a mean variance trend, allowing better modelling of low abundance genes. Finally, test of differential gene expression is carried out by testing the significance of one or more coefficients of the fitted linear model.

### Wilcoxon Rank Sum (WRS) Test

WRS^34^ test is a non-parametric test commonly used for pairwise OGE testing. The test is formulated as *H*_0_, the distributions of the gene vector under two dose groups are equal versus *H*_*a*_: the distributions are not equal.The test involves the calculation of the *U* statistic, which for large samples is approximtaly normally distributed. Since this is a pairwise test, a union is taken over all the genes found to be DE in each of the pairwise tests. The computations are carried out using the *wilcox*.*test* function in R package *stats* and multiplicity is controlled using FDR correction.

### ANOVA

Analysis of variance (ANOVA) ^29^ is very commonly used for testing the differences among means in multiple groups. For a fixed gene *j*, it is assumed that the observed gene vector *y*_*k,i,j*,_ for cell *i* is grouped by dose. Assuming that 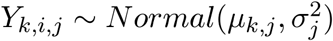, ANOVA aims to test the null hypothesis *H*_0_: *μ*_1,*j*_ = *μ*_2,*j*_ = … *μ*_*K,j*_ = *μ*_*j*_ versus *H*_*a*_ : *μ*_*k,j*_, *i* = 1, …, *n;j* = 1, …, *p; k* = 1, …, *K* is different for at least one *k*. The test statistic is computed using the *aov* function in R package *stats* and it follows a F-distribution with *(K* − 1) and *(n* − *K)* degrees of freedom. Multiplicity is controlled by applying FDR correction on the obtained p-values.

### Kruskal-Wallis (KW) Test

KW^30^ test extends the WRS test for multiple groups. It is also a non-parametric extension of the ANOVA test.The test is formulated as; *H*_0_, the distributions of the gene vector under K dose groups are equal versus *H*_*a*_: the distributions are not equal. The computation of the KW test statistic is carried out using the *kruskal*.*test* function in R- package *stats* and it asymptotically follows a *χ*^2^ distribution with *K* − 1 degrees of freedom. Multiplicity is controlled by applying FDR correction on the obtained p-values.

### Benchmarking and sensitivity analyses

Benchmarking of DE test methods was performed on simulated datasets based on initial parameters derived from real dose-response snRNAseq data. The probability of differential expression was set to 10% with a 50% probability of being down-regulated, equally distributed among the dose-response models in Table 1. Batch parameters were used to include sample variation associated with data obtained from 3 individuals in each dose group. A total of 5,000 genes were simulated for 4,500 cells (500 per dose group) using the same doses as the real dataset. Sensitivity analyses varied each of the following parameters according to values is supplementary Table 1: cell abundance equally distributed among dose groups, varying cell numbers in each dose group, percent DE genes, proportion of downregulated DE genes, fold-change location or scale, and dropout rate. Each simulation was replicated 10 times using a different initial seed. Method concordance was determined as area under the concordance curve (AUCC) for the top 100- or 500-ranked genes in simulated and real datasets, respectively, as previously described^37^.

## Acknowledgements

This work was supported by the National Human Genome Research Institute R21 HG010789 to T.R. Zacharewski and S. Bhattacharya, and the National Institutes of Environmental Health Sciences Superfund Research Program P42 ES004911 to T.R. Zacharewski. T.R. Zacharewski and S. Bhattacharya are partially supported by AgBioResearch at Michigan State University. T. Maiti and S. Saha are partially supported by NSF-DMS 1945824.

## Author contributions

R. Nault and T.R. Zacharewski designed the in-life study. R. Nault performed the *in vivo* experiments and generated the TCDD dose-response experimental dataset. R. Nault and S. Saha performed analyses, data interpretation, and prepared the first manuscript draft. S. Saha and S. Sinha developed statistical tests. R .Nault, S. Saha., and J. Dodson contributed to code development. S. Bhattacharya, S. Sinha, T. Maiti, and T.R. Zacharewski supported the analyses and interpretation of outcomes. All authors contributed and reviewed the manuscript.

## Ethics declarations

The authors have no competing interests to declare.

---

**Supplementary Figure 1:**
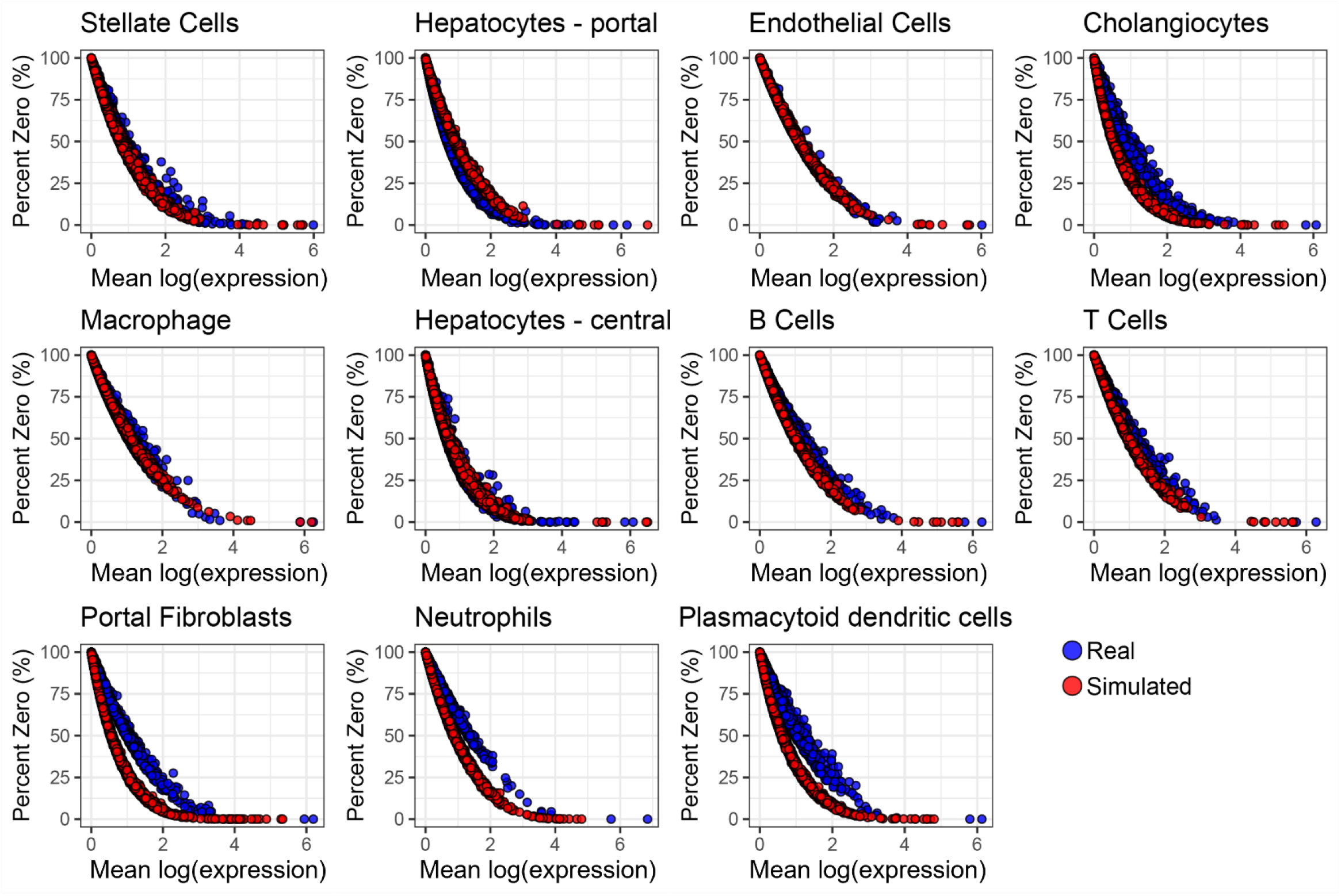
Comparison of experimental and simulated dose-response data from cell specific initial parameters. Simulation parameters were estimated from each cell type identified in our experimental hepatic dose-response snRNAseq dataset. A total of 4500 cells (500 per dose) distributed across 3 individuals for 9 dose groups were simulated. The percent zeroes and mean log expression was calculated for each gene for all dose groups combined.

**Supplementary Figure 2:**
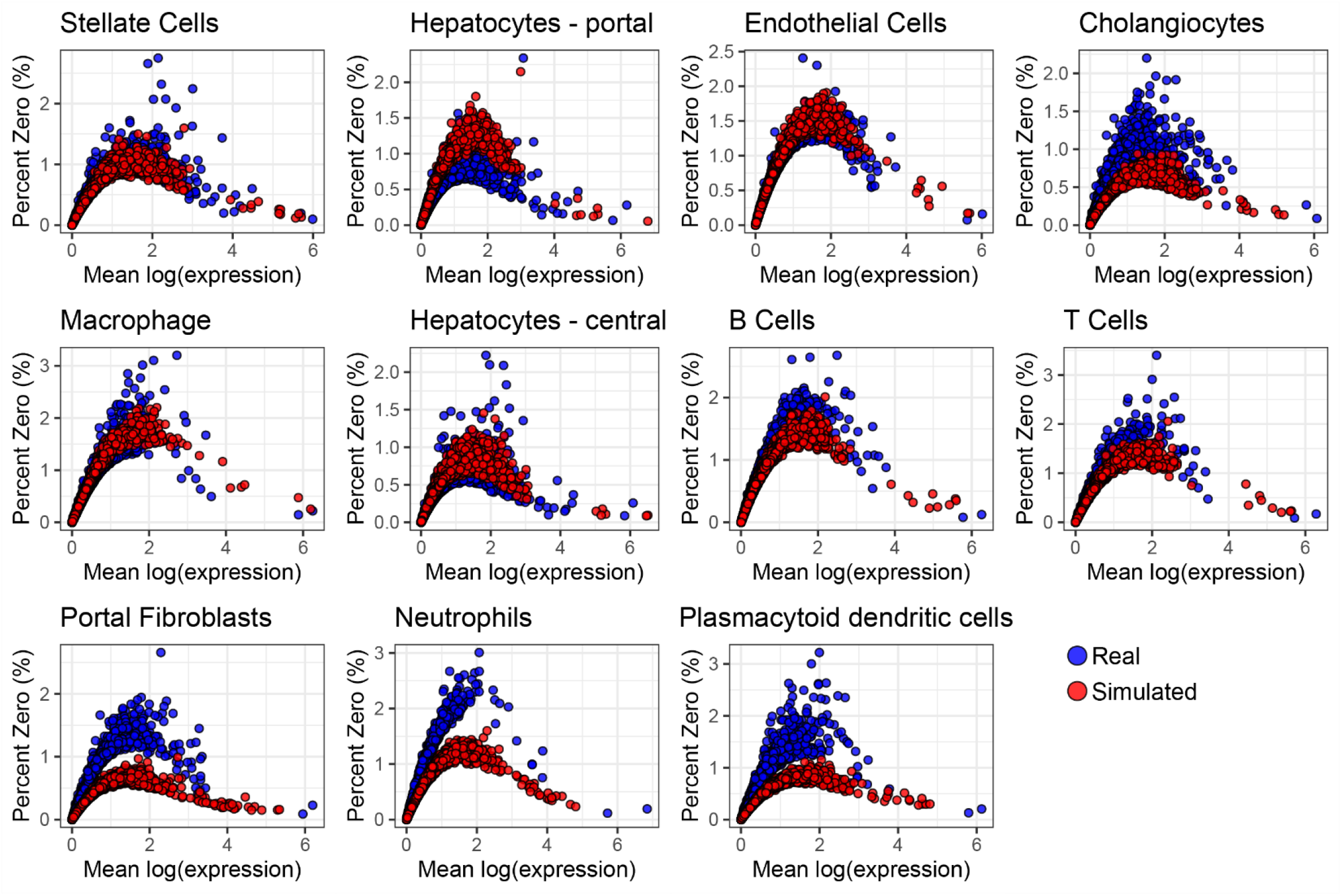
Comparison of experimental and simulated dose-response data from cell specific initial parameters. Simulation parameters were estimated from each cell type identified in our experimental hepatic dose-response snRNAseq dataset. A total of 4500 cells (500 per dose) distributed across 3 individuals for 9 dose groups were simulated. The percent zeroes and mean log variance was calculated for each gene for all dose groups combined.

**Supplementary Figure 3:**
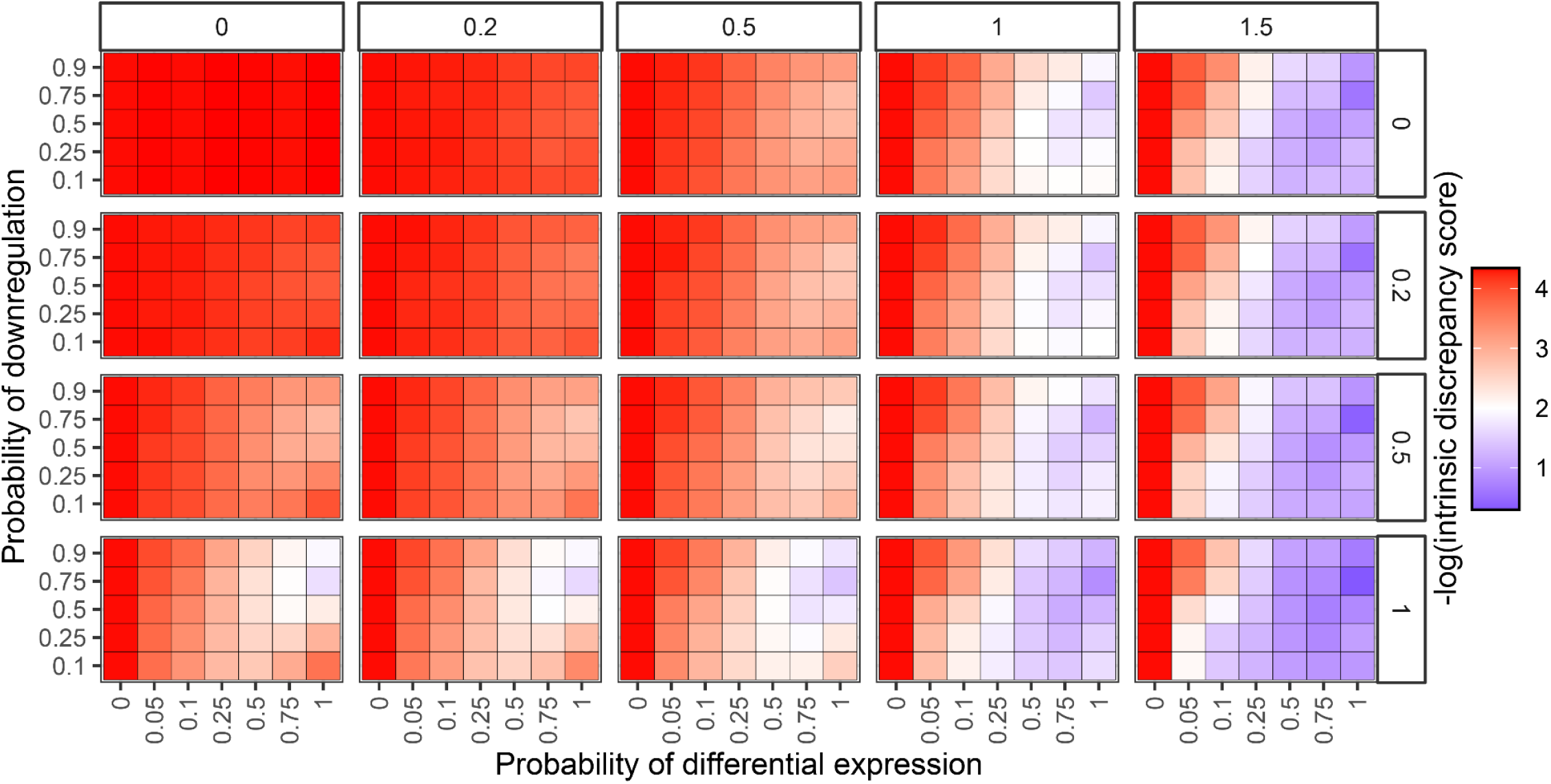
Intrinsic discrepancy scores of fold-change distributions under varying simulation parameters. Datasets were simulated for 5,000 genes by varying the probability of differential expression, probability of repression, mean fold-change of differentially expressed genes (location) and distribution of fold-change for differentially expressed genes (scale). Fold-change location and scale of differentially expressed genes from 0 – 1.5 and 0 – 1, respectively, represent the values for mean and standard deviation of a log-normal distribution. 5,000 simulated genes were compared to an equivalent number sampled from experimental data for the fold-changes between 30 µg/kg TCDD and control groups. The Kullback-Leibler Divergence (KLD) intrinsic discrepancy (ID) was used to evaluate the similarity in distributions.

**Supplementary Figure 4:**
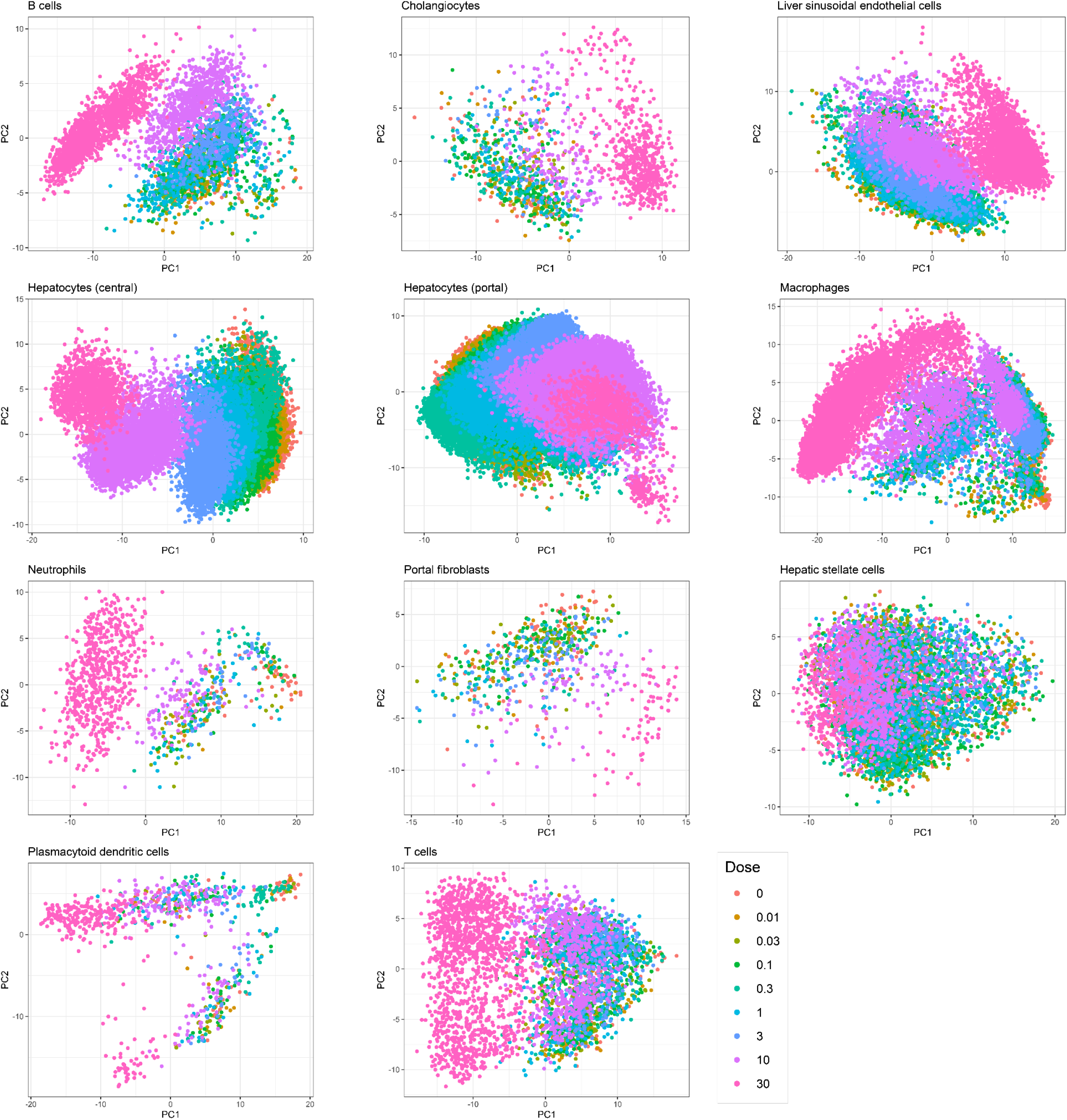
Principal components analysis (PCA) of experimental hepatic dose-response data from male mice gavaged with 2,3,7,8-tetrachlorodibenzo-p-dioxin (TCDD) every 4 days for 28 days. PCA was performed for all genes and each identified cell type. Each point represents an individual cell. Colors represent treatment groups.

**Supplementary Figure 5:**
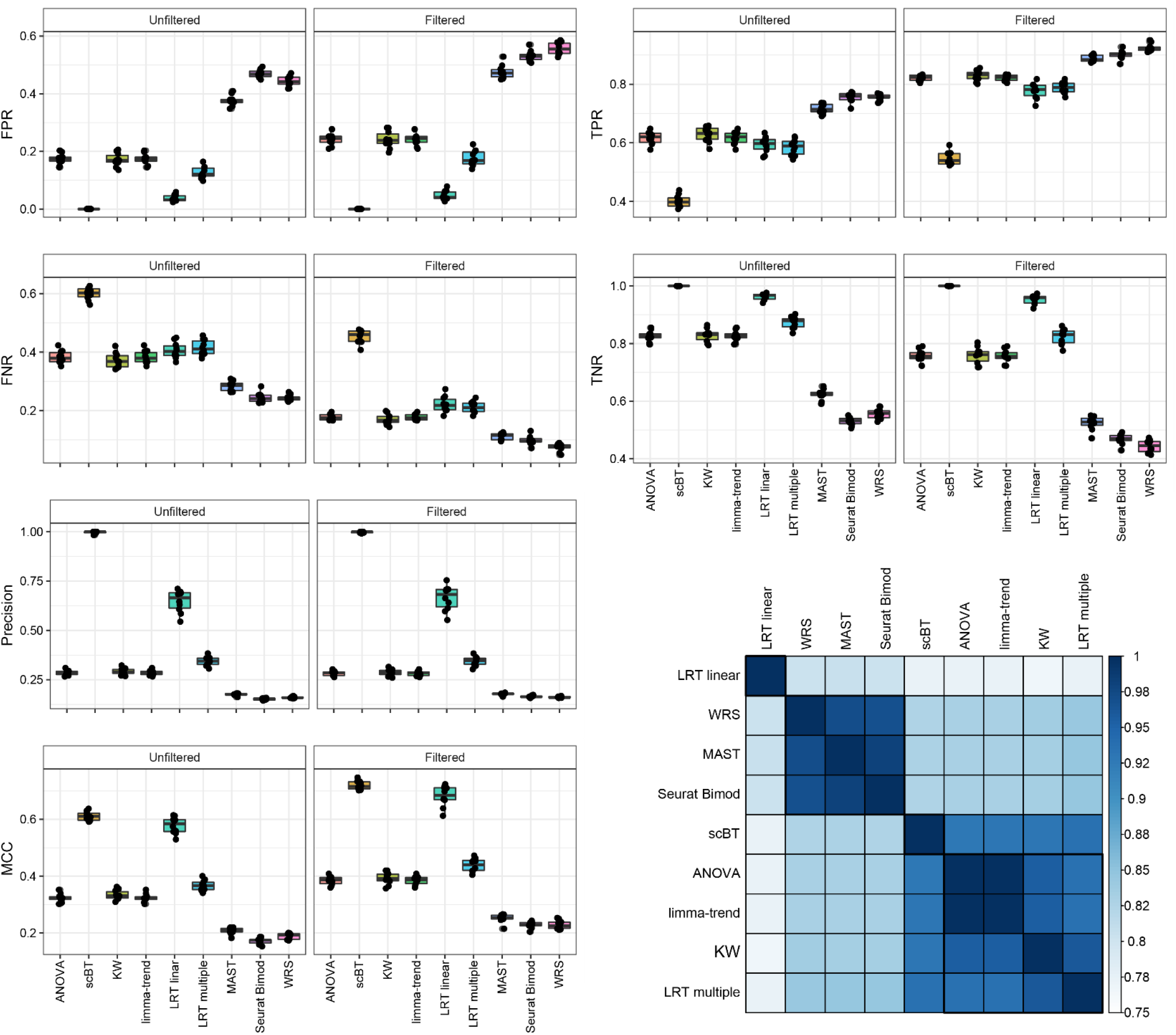
Benchmarking metrics of 9 differential expression test methods for simulated dose response data using default initial parameters. A total of 4,500 cells (500 cells per dose group) and 5,000 genes were simulated across 9 dose groups with a probability of being differentially expressed of 10%, of which 50% were repressed. Differential expression fold-change location and scale were 0.8 and 0.4, respectively. Given a ground truth from simulation outputs, false positive rates (FPR), true positive rates (TPR), false negative rates (FNR), true negative rates (TNR), precision, and Matthews correlation coefficient (MCC) were calculated. Points represent median ± minimum to maximum values for 10 replicate simulations. Heat map represents the area under the concordance curve (AUCC) of the 100 most significant gene expression changes (lowest *P*-values) calculated for each pairwise comparison and clustered based on the similarity of the scores. Box and whisker plots represent median and 25^th^ and 75^th^ percentile, and minimum and maximum values for 10 replicate simulations.

**Supplementary Figure 6:**
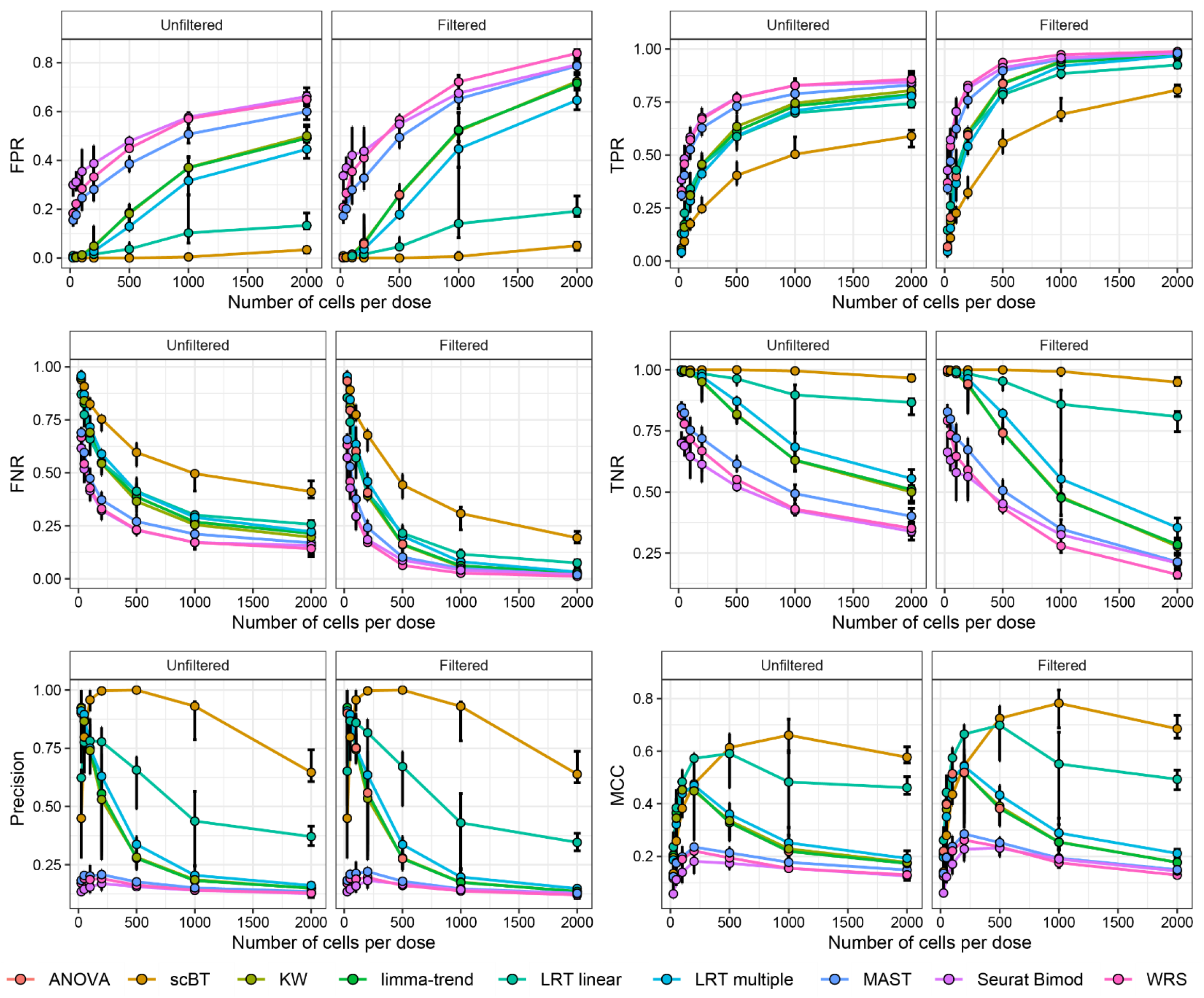
Benchmarking metrics of 9 differential expression test methods for simulated dose response data with varying cell abundances. 5,000 genes were simulated across 9 dose groups with a probability of being differentially expressed of 10%, of which 50% were repressed. Differential expression fold-change location and scale were 0.8 and 0.4, respectively. Given a ground truth from simulation outputs, false positive rates (FPR), true positive rates (TPR), false negative rates (FNR), true negative rates (TNR), precision, and Matthews correlation coefficient (MCC) were calculated. Points represent median ± minimum to maximum values for 10 replicate simulations.

**Supplementary Figure 7:**
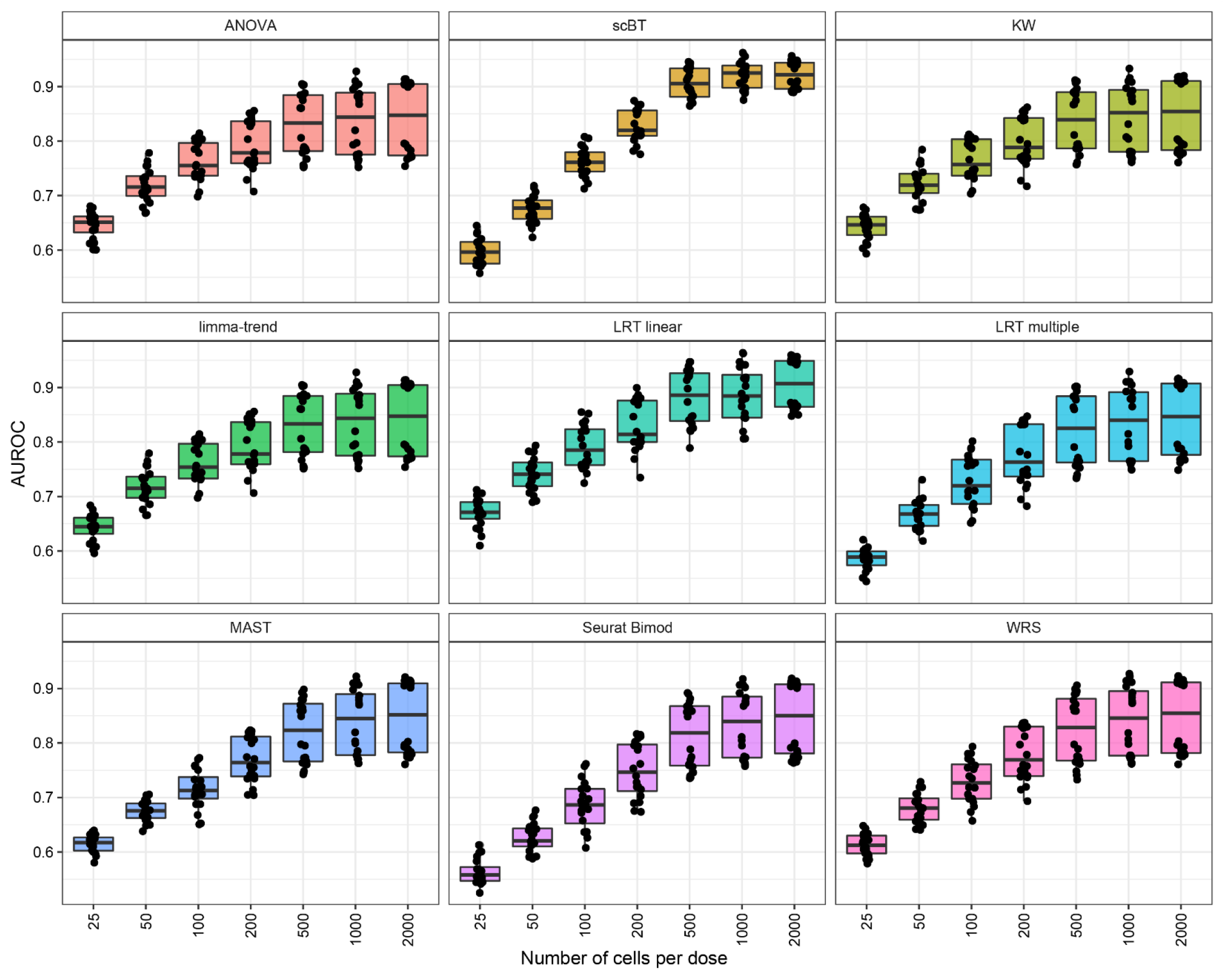
Area under the receiver-operating curve (AUROC) of 9 differential expression test methods for simulated dose response data with varying cell abundances. 5,000 genes were simulated across 9 dose groups with a probability of being differentially expressed of 10%, of which 50% were repressed. Differential expression fold-change location and scale were 0.8 and 0.4, respectively. Box and whisker plots represent median and 25^th^ and 75^th^ percentile, and minimum and maximum values for 10 replicate simulations.

**Supplementary Figure 8:**
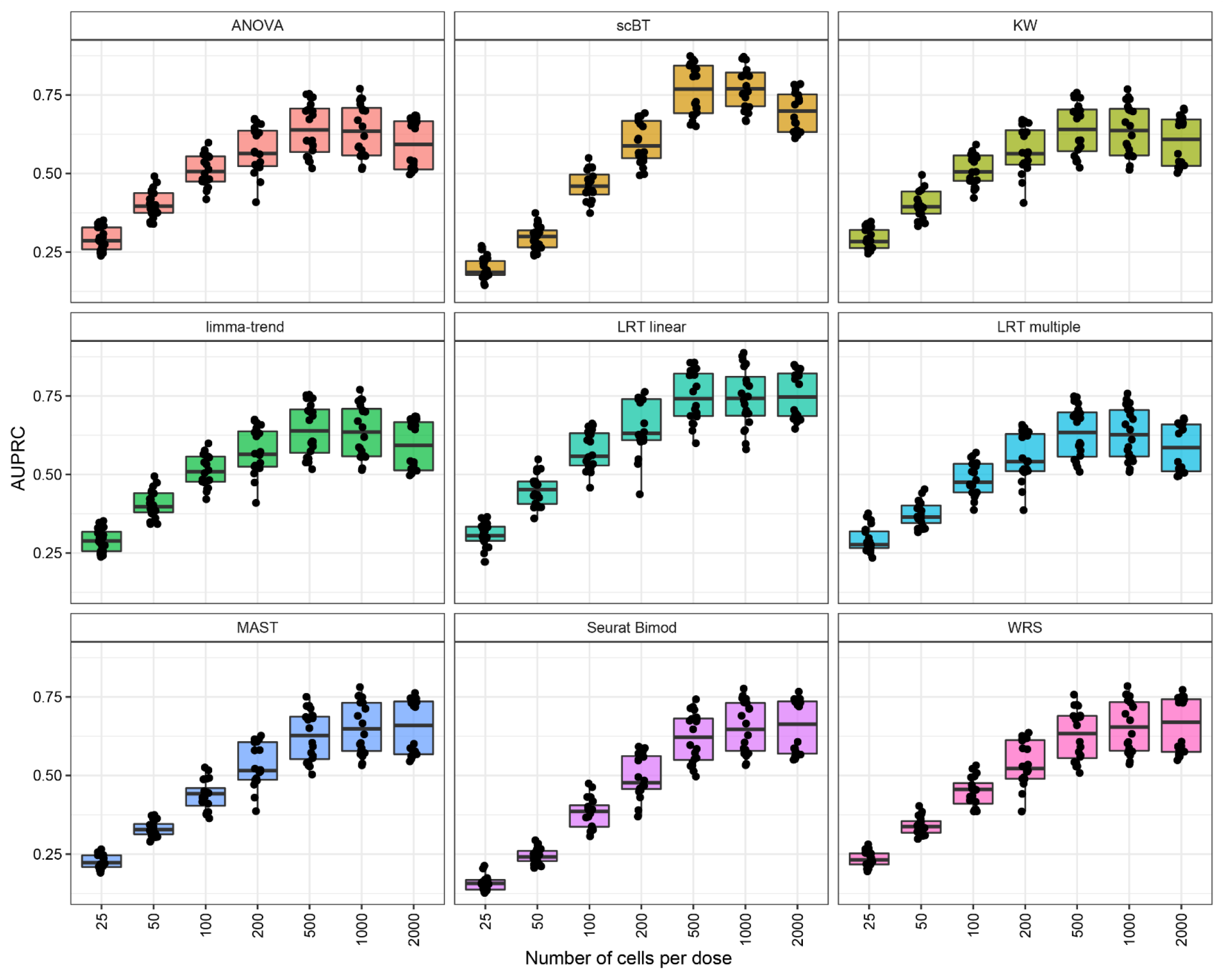
Area under the precision-recall curve (AUPRC) of 9 differential expression test methods for simulated dose response data with varying number of cell abundances. 5,000 genes were simulated across 9 dose groups with a probability of being differentially expressed of 10%, of which 50% were repressed. Differential expression fold-change location and scale were 0.8 and 0.4, respectively. Box and whisker plots represent median and 25^th^ and 75^th^ percentile, and minimum and maximum values for 10 replicate simulations.

**Supplementary Figure 9:**
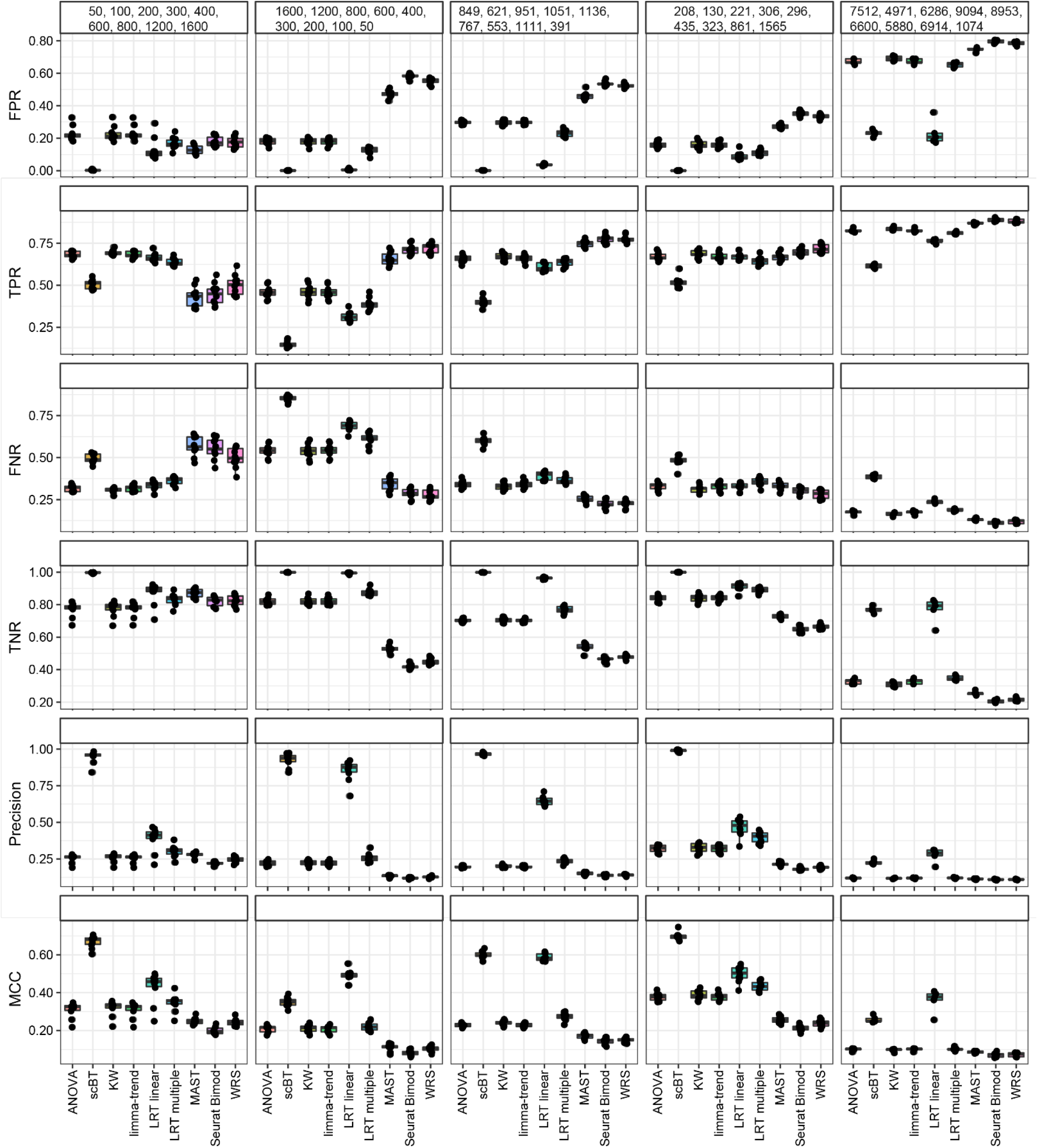
Benchmarking metrics of 9 differential expression test methods for simulated dose response data with varying number of cells per dose group. 5,000 genes were simulated across 9 dose groups with a probability of being differentially expressed of 10%, of which 50% were repressed. Differential expression fold-change location and scale were 0.8 and 0.4, respectively. Given a ground truth from simulation outputs, false positive rates (FPR), true positive rates (TPR), false negative rates (FNR), true negative rates (TNR), precision, and Matthews correlation coefficient (MCC) were calculated. Box and whisker plots represent median and 25^th^ and 75^th^ percentile, and minimum and maximum values for 10 replicate simulations.

**Supplementary Figure 10:**
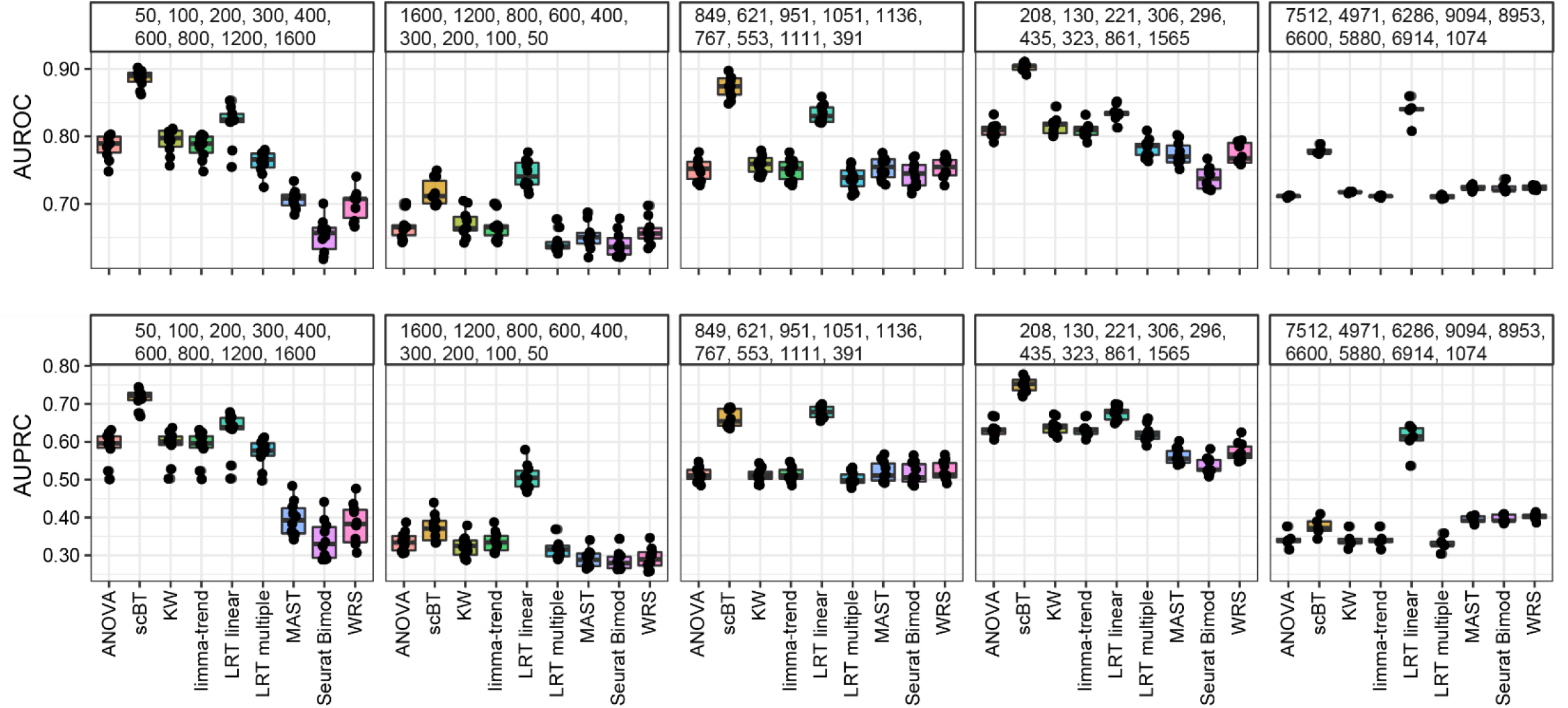
Area under the receiver-operating curve (AUROC) and area under the precision-recall curve (AUPRC) of 9 differential expression test methods for simulated dose response data with varying number of cells per dose group. 5,000 genes were simulated across 9 dose groups with a probability of being differentially expressed of 10%, of which 50% were repressed. Box and whisker plots represent median and 25^th^ and 75^th^ percentile, and minimum and maximum values for 10 replicate simulations.

**Supplementary Figure 11:**
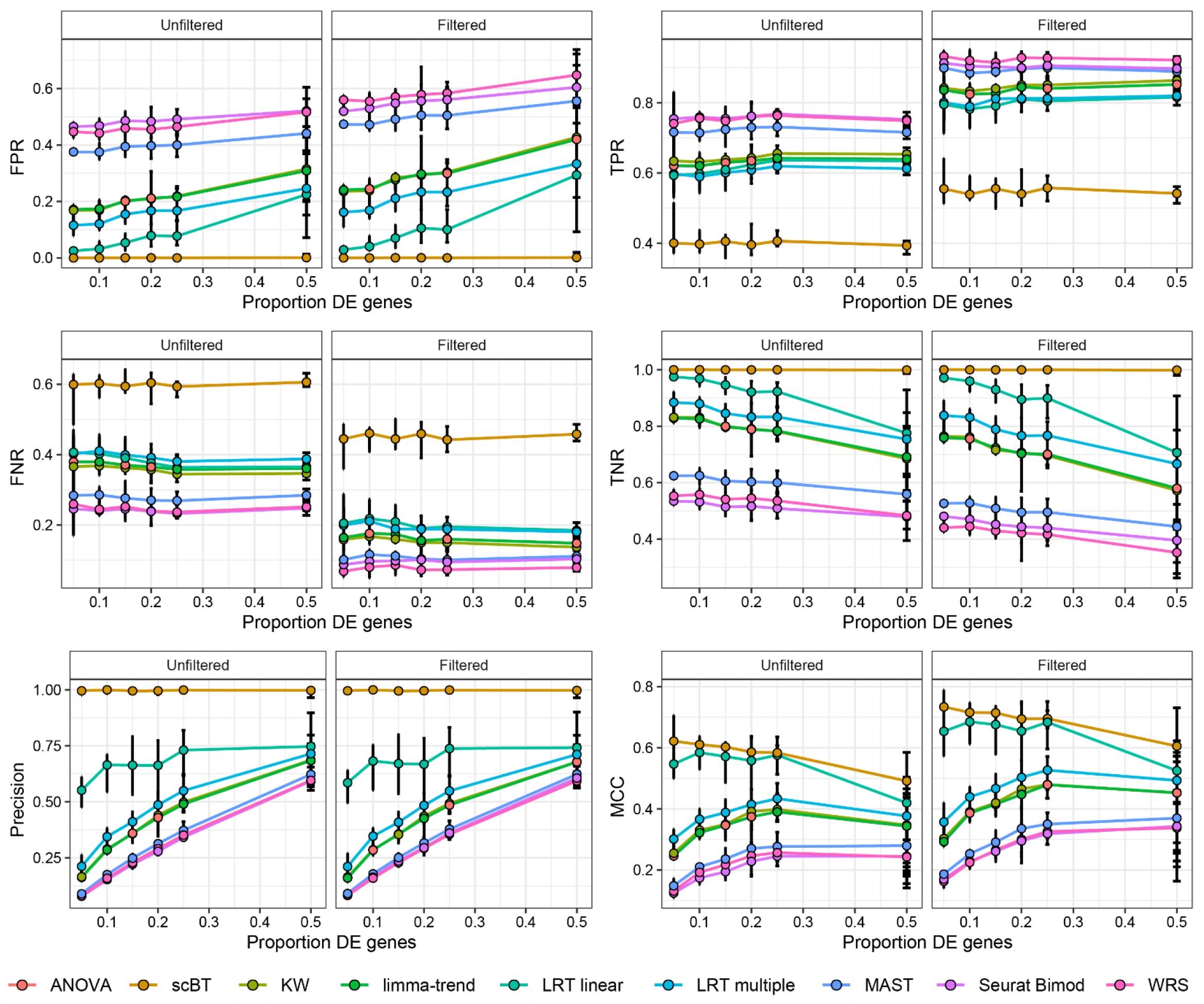
Benchmarking metrics of 9 differential expression test methods for simulated dose response data with varying differential expression probabilities. A total of 4,500 cells (500 cells per group) and 5,000 genes were simulated across 9 dose groups with a 50% probability of being repressed. Differential expression fold-change location and scale were 0.8 and 0.4, respectively. Given a ground truth from simulation outputs, false positive rates (FPR), true positive rates (TPR), false negative rates (FNR), true negative rates (TNR), precision, and Matthews correlation coefficient (MCC) were calculated. Points represent median ± minimum to maximum values for 10 replicate simulations.

**Supplementary Figure 12:**
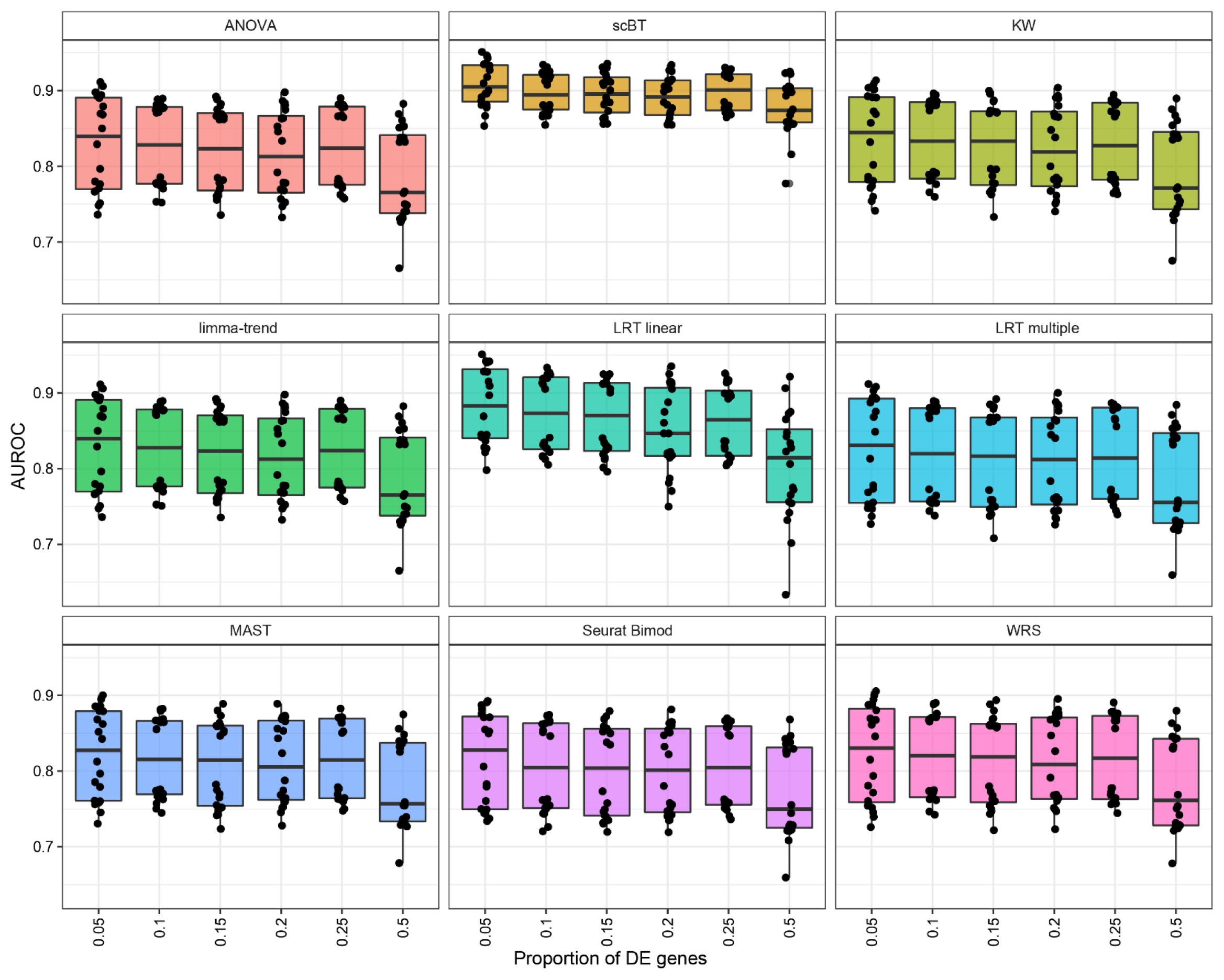
Area under the receiver-operating curve (AUROC) of 9 differential expression test methods for simulated dose response data with varying differential expression probabilities. A total of 4,500 cells (500 cells per group) and 5,000 genes were simulated across 9 dose groups with a 50% probability of being repressed. Differential expression fold-change location and scale were 0.8 and 0.4, respectively. Box and whisker plots represent median and 25^th^ and 75^th^ percentile, and minimum and maximum values for 10 replicate simulations.

**Supplementary Figure 13:**
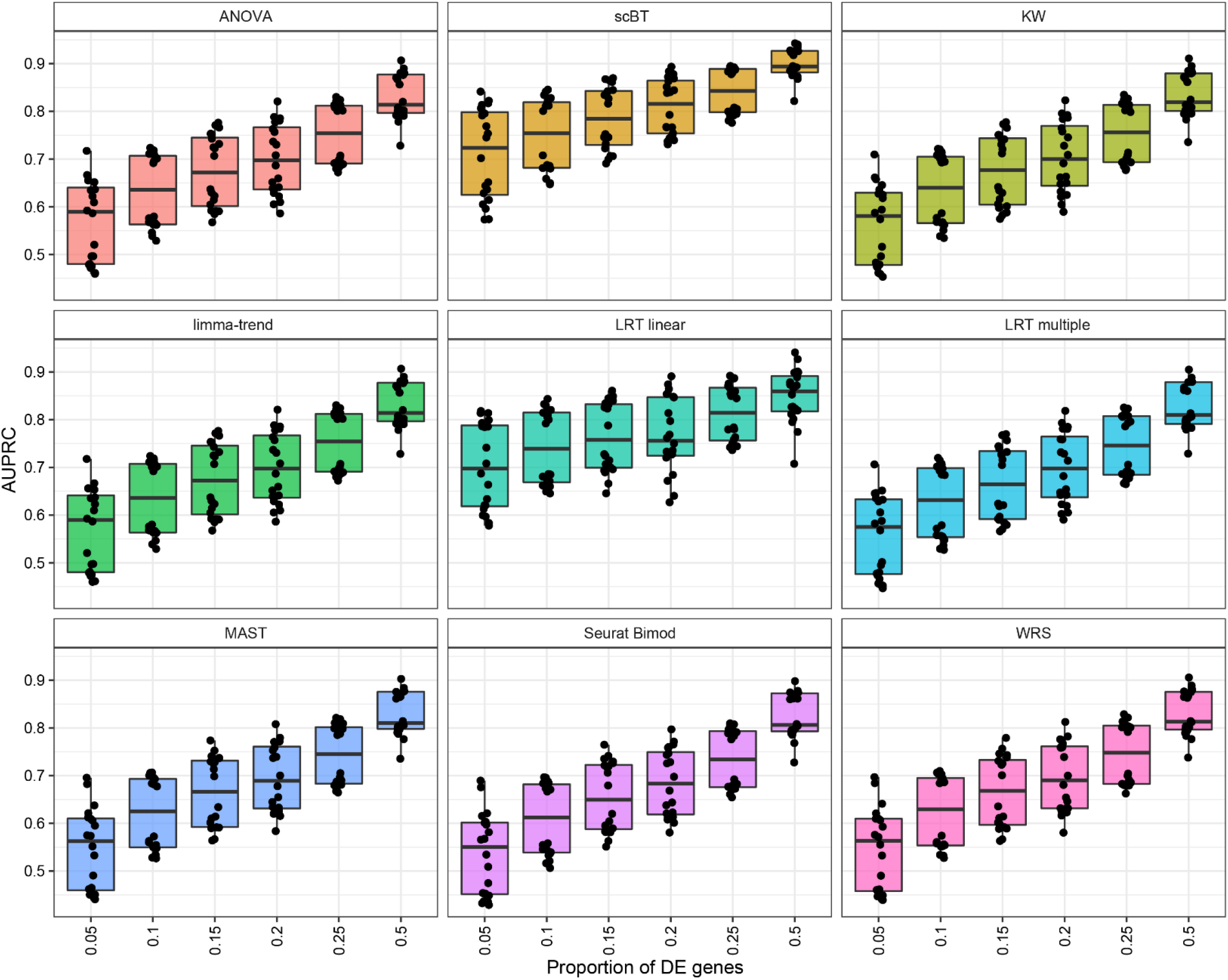
Area under the receiver-operating curve (AUPRC) of 9 differential expression test methods for simulated dose response data with varying differential expression probabilities. A total of 4,500 cells (500 cells per group) and 5,000 genes were simulated across 9 dose groups with a 50% probability of being repressed. Differential expression fold-change location and scale were 0.8 and 0.4, respectively. Box and whisker plots represent median and 25^th^ and 75^th^ percentile, and minimum and maximum values for 10 replicate simulations.

**Supplementary Figure 14:**
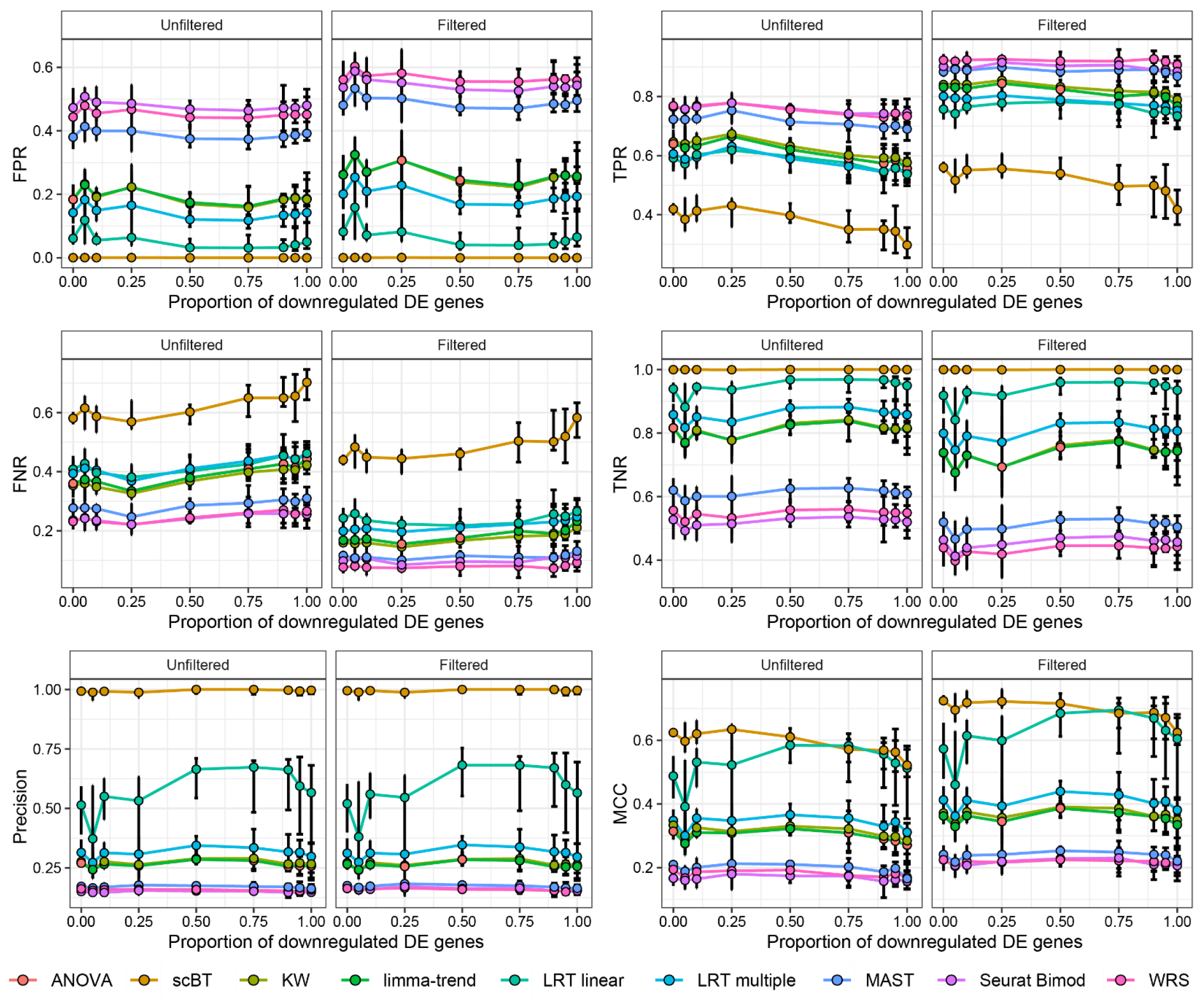
Benchmarking metrics of 9 differential expression test methods for simulated dose response data with varying probability of repressed genes. A total of 4,500 cells (500 cells per group) and 5,000 genes were simulated across 9 dose groups with a 10% probability of differential expression. Differential expression fold-change location and scale were 0.8 and 0.4, respectively. Given a ground truth from simulation outputs, false positive rates (FPR), true positive rates (TPR), false negative rates (FNR), true negative rates (TNR), precision, and Matthews correlation coefficient (MCC) were calculated. Points represent median ± minimum to maximum values for 10 replicate simulations.

**Supplementary Figure 15:**
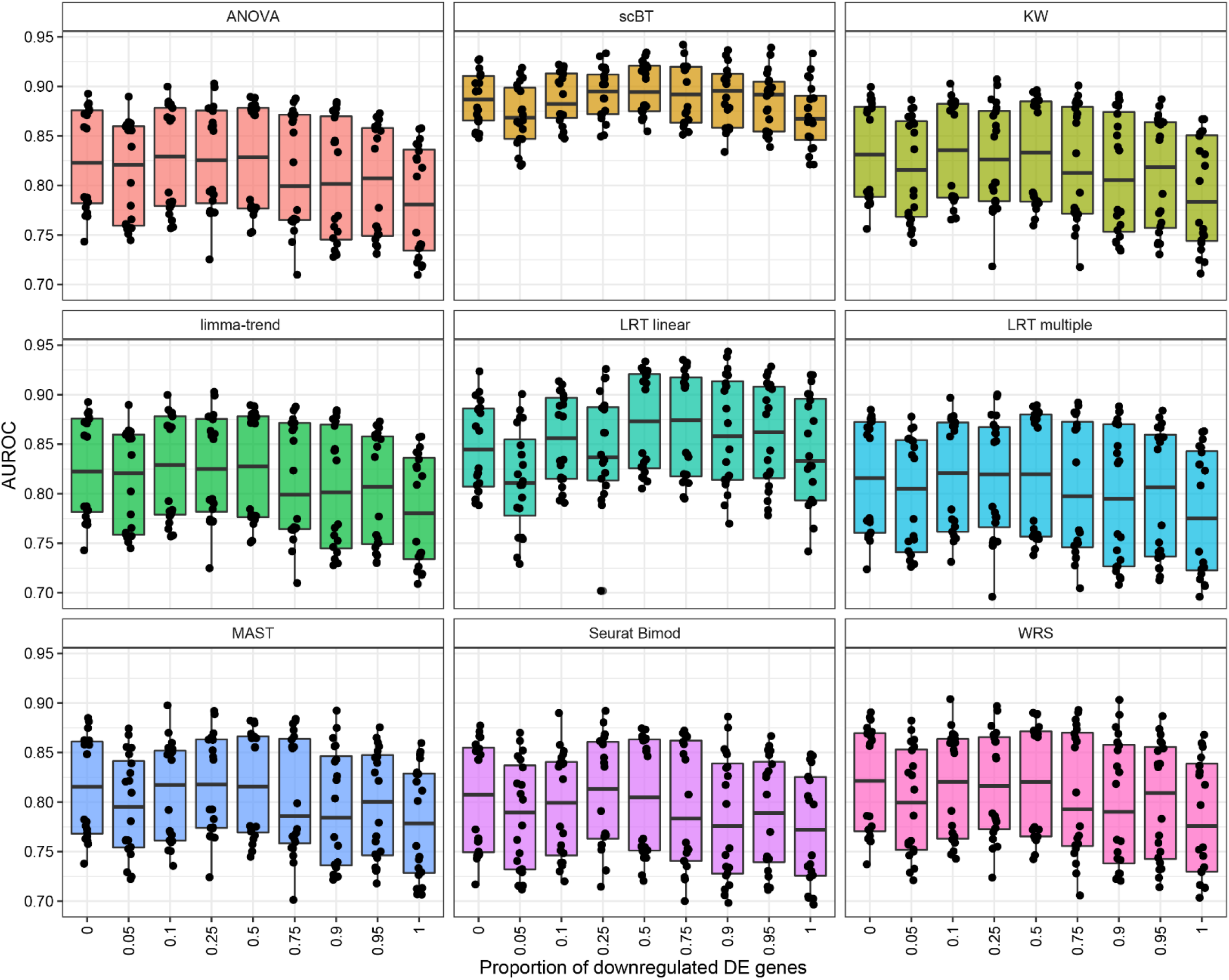
Area under the receiver-operating curve (AUROC) of 9 differential expression test methods for simulated dose response data with varying probability of repressed genes. A total of 4,500 cells (500 cells per group) and 5,000 genes were simulated across 9 dose groups with a 10% probability of differential expression. Differential expression fold-change location and scale were 0.8 and 0.4, respectively. Box and whisker plots represent median and 25^th^ and 75^th^ percentile, and minimum and maximum values for 10 replicate simulations.

**Supplementary Figure 16:**
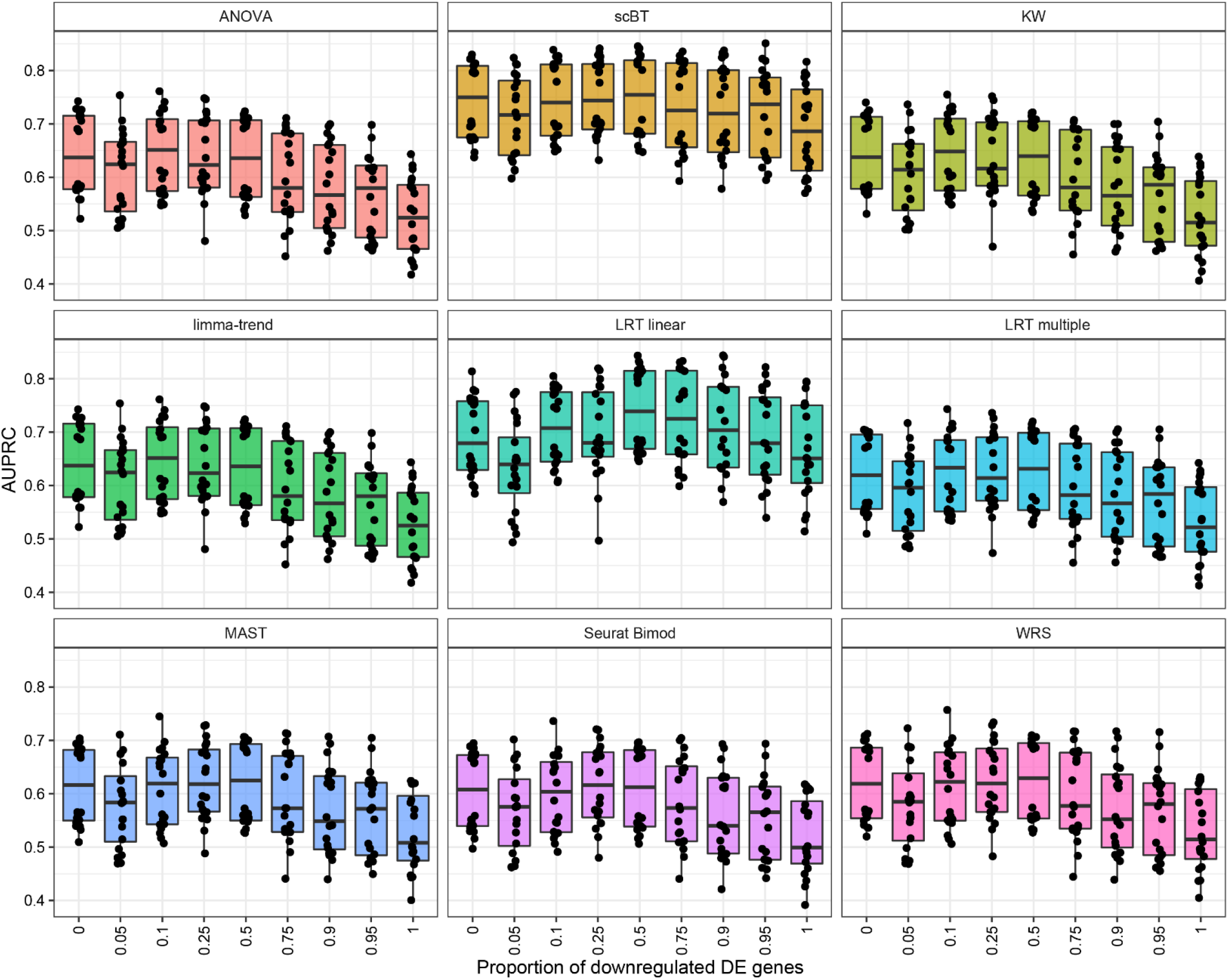
Area under the receiver-operating curve (AUPRC) of 9 differential expression test methods for simulated dose response data with varying probability of repressed genes. A total of 4,500 cells (500 cells per group) and 5,000 genes were simulated across 9 dose groups with a 10% probability of differential expression. Differential expression fold-change location and scale were 0.8 and 0.4, respectively. Box and whisker plots represent median and 25^th^ and 75^th^ percentile, and minimum and maximum values for 10 replicate simulations.

**Supplementary Figure 17:**
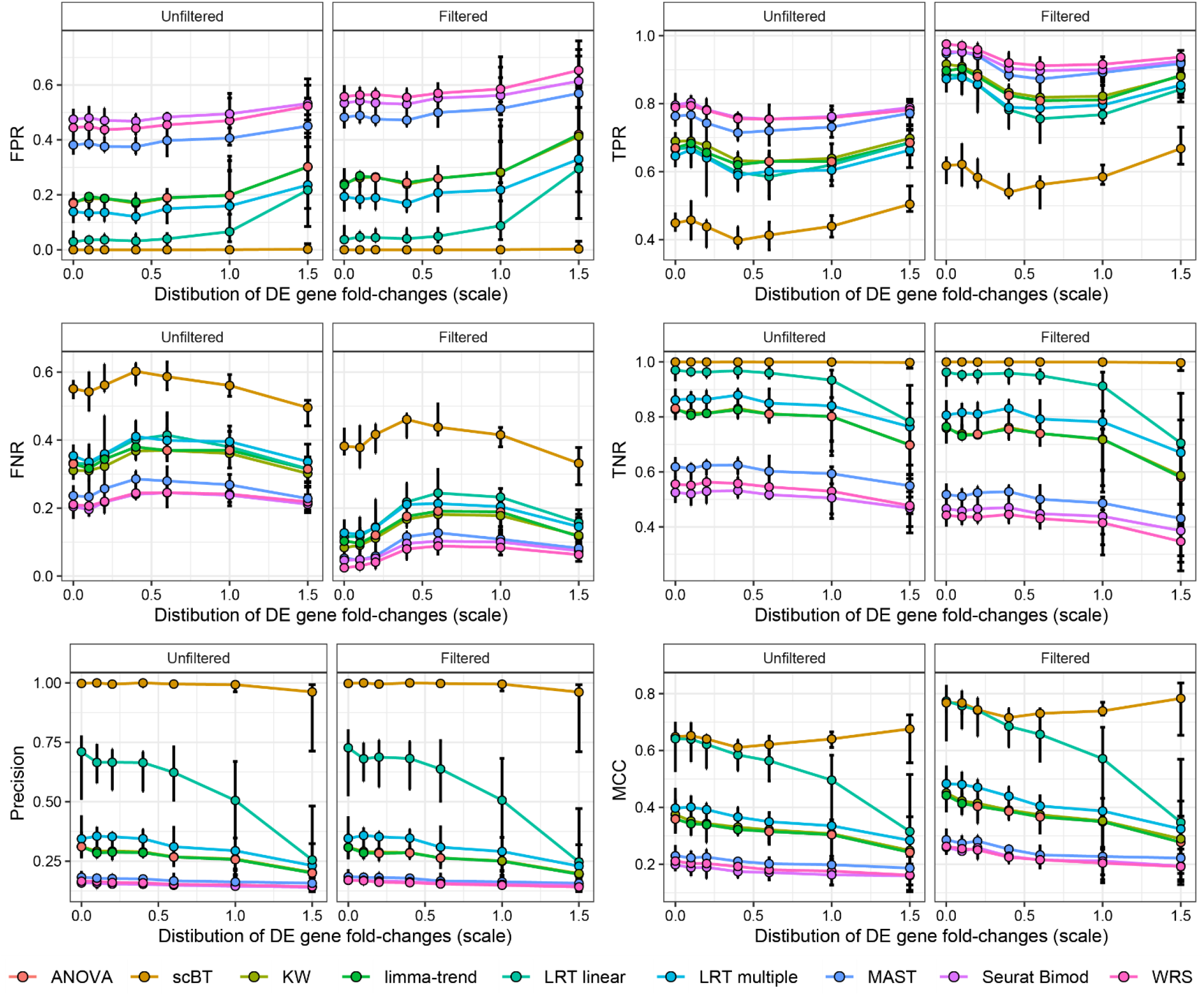
Benchmarking metrics of 9 differential expression test methods for simulated dose response data with varying scale of differentially expressed genes. A total of 4,500 cells (500 cells per group) and 5,000 genes were simulated across 9 dose groups with a probability of being differentially expressed of 10%, of which 50% were repressed. Differential expression fold-change location was 0.8. Given a ground truth from simulation outputs, false positive rates (FPR), true positive rates (TPR), false negative rates (FNR), true negative rates (TNR), precision, and Matthews correlation coefficient (MCC) were calculated. Points represent median ± minimum to maximum values for 10 replicate simulations.

**Supplementary Figure 18:**
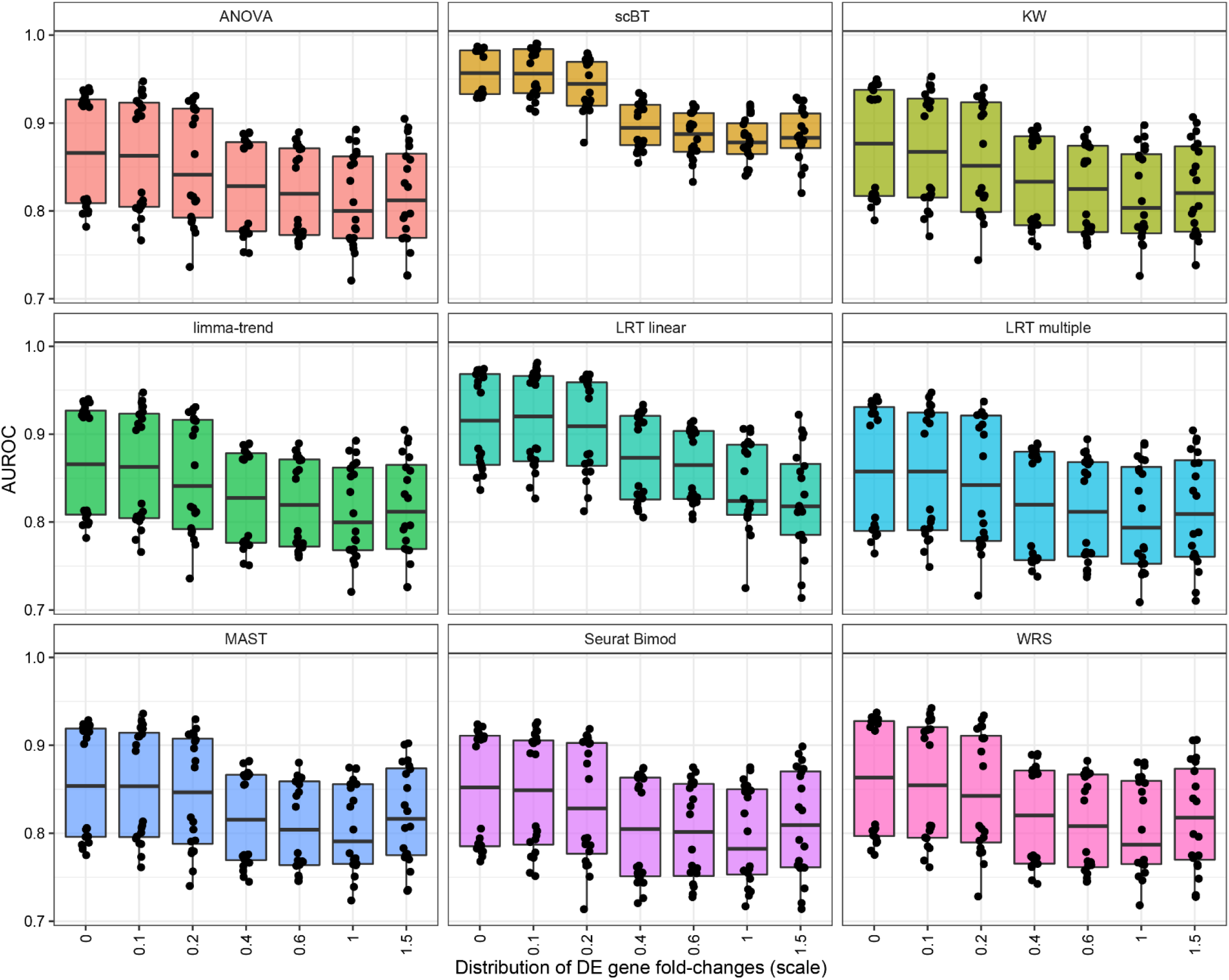
Area under the receiver-operating curve (AUROC) of 9 differential expression test methods for simulated dose response data with varying scale of differentially expressed genes. A total of 4,500 cells (500 cells per group) and 5,000 genes were simulated across 9 dose groups with a probability of being differentially expressed of 10%, 50% of which were repressed. Differential expression fold-change location was 0.8. Box and whisker plots represent median and 25^th^ and 75^th^ percentile, and minimum and maximum values for 10 replicate simulations.

**Supplementary Figure 19:**
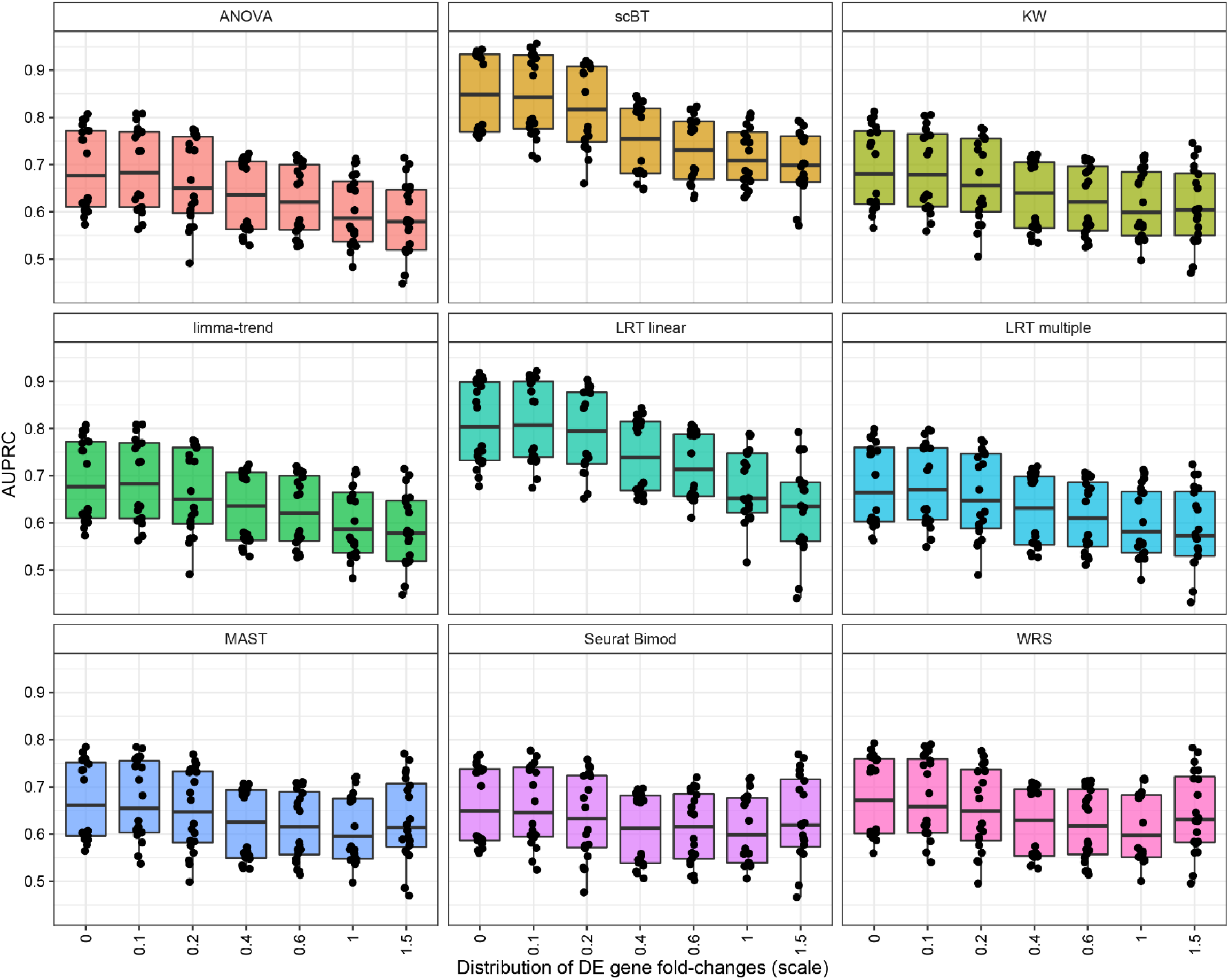
Area under the precision-recall curve (AUPRC) of 9 differential expression test methods for simulated dose response data with varying scale of differentially expressed genes. A total of 4,500 cells (500 cells per group) and 5,000 genes were simulated across 9 dose groups with a probability of being differentially expressed of 10%, of which 50% were repressed. Differential expression fold-change location was 0.8. Box and whisker plots represent median and 25^th^ and 75^th^ percentile, and minimum and maximum values for 10 replicate simulations.

**Supplementary Figure 20:**
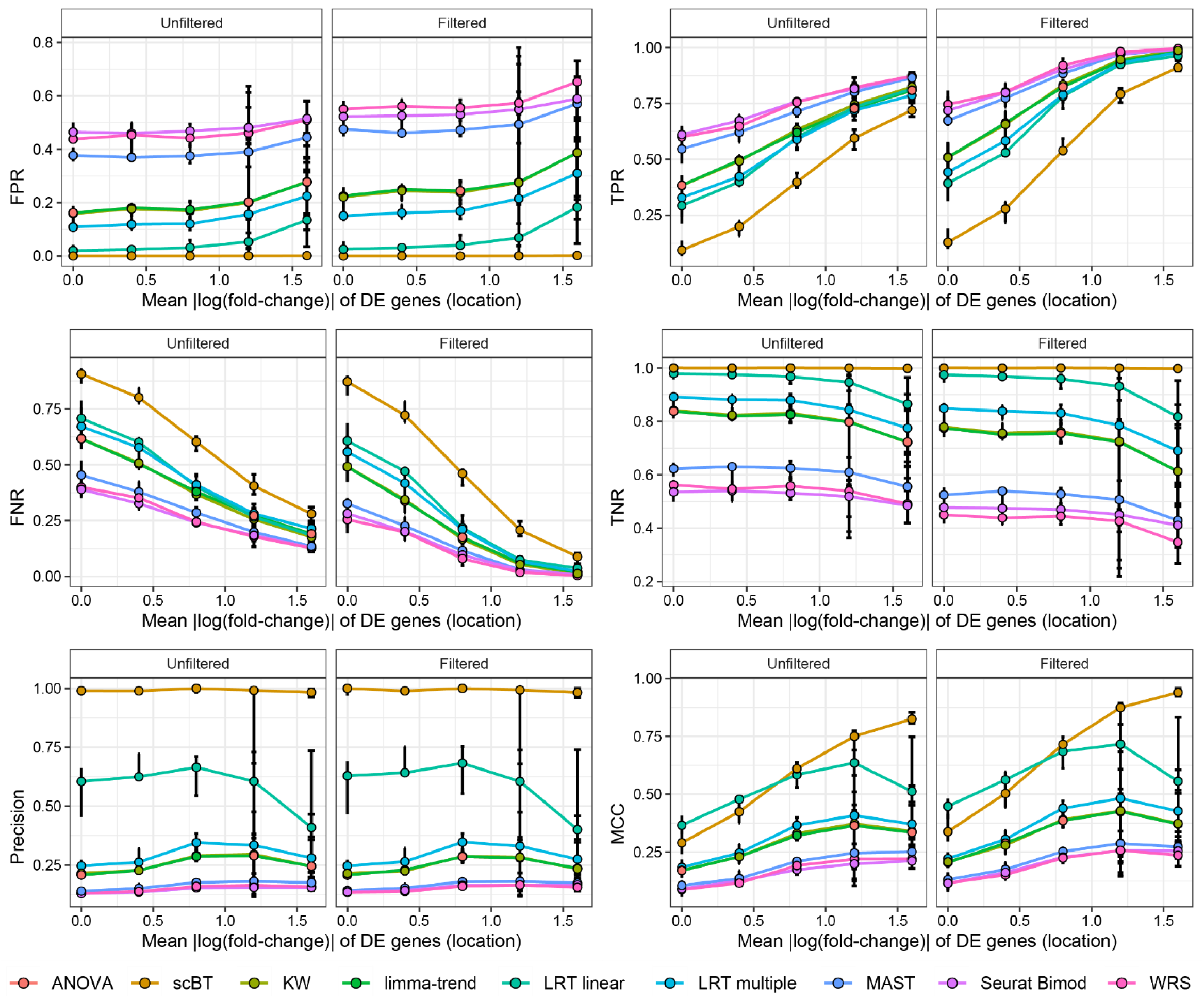
Benchmarking metrics of 9 differential expression test methods for simulated dose response data with varying location of differentially expressed genes. A total of 4,500 cells (500 cells per group) and 5,000 genes were simulated across 9 dose groups with a probability of being differentially expressed of 10%, of which 50% were repressed. Differential expression fold-change scale was 0.4. Given a ground truth from simulation outputs, false positive rates (FPR), true positive rates (TPR), false negative rates (FNR), true negative rates (TNR), precision, and Matthews correlation coefficient (MCC) were calculated. Points represent median ± minimum to maximum values for 10 replicate simulations.

**Supplementary Figure 21:**
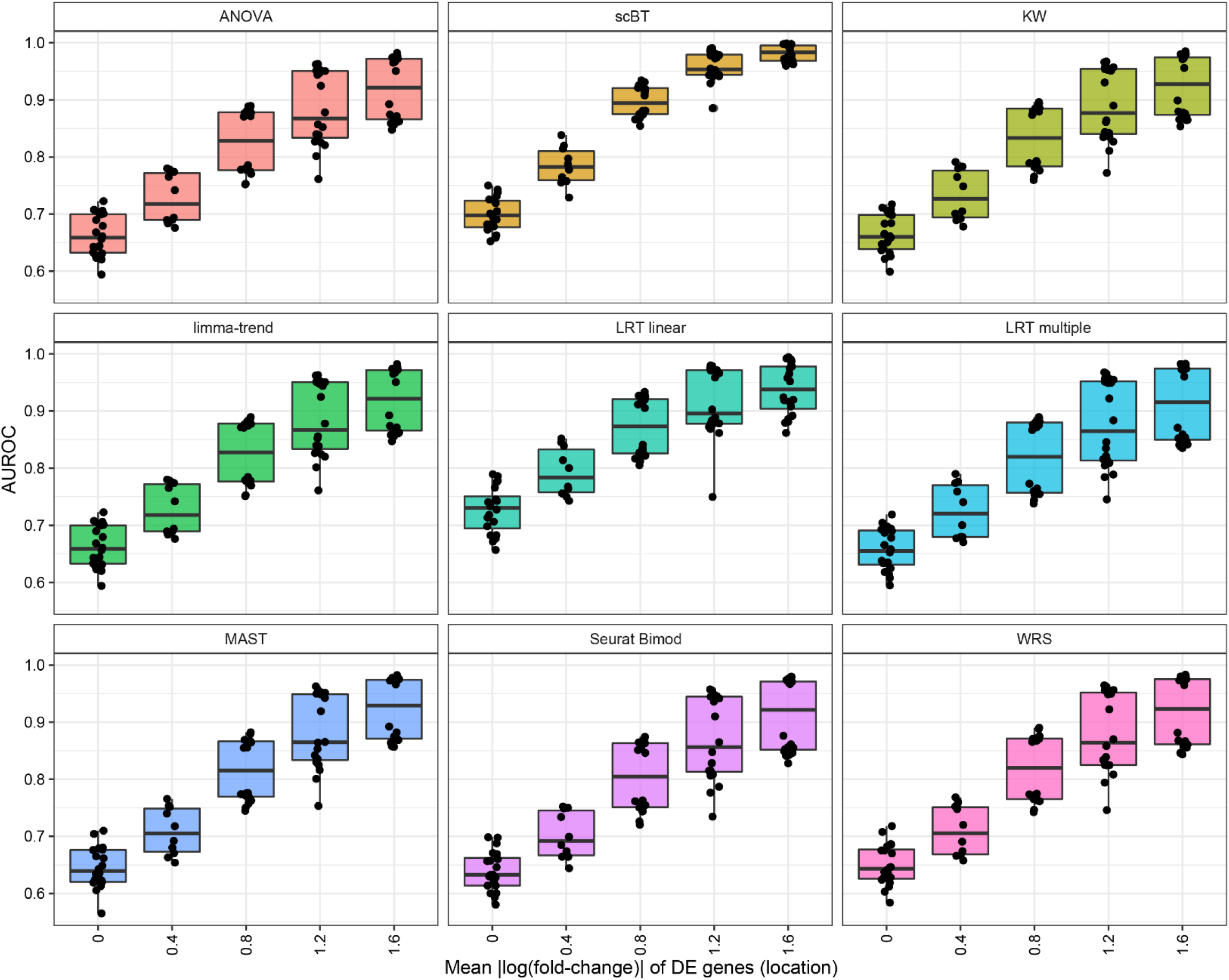
Area under the receiver-operating curve (AUROC) of 9 differential expression test methods for simulated dose response data with varying location of differentially expressed genes. A total of 4,500 cells (500 cells per group) and 5,000 genes were simulated across 9 dose groups with a probability of being differentially expressed of 10%, of which 50% were repressed. Differential expression fold-change scale was 0.4. Box and whisker plots represent median and 25^th^ and 75^th^ percentile, and minimum and maximum values for 10 replicate simulations.

**Supplementary Figure 22:**
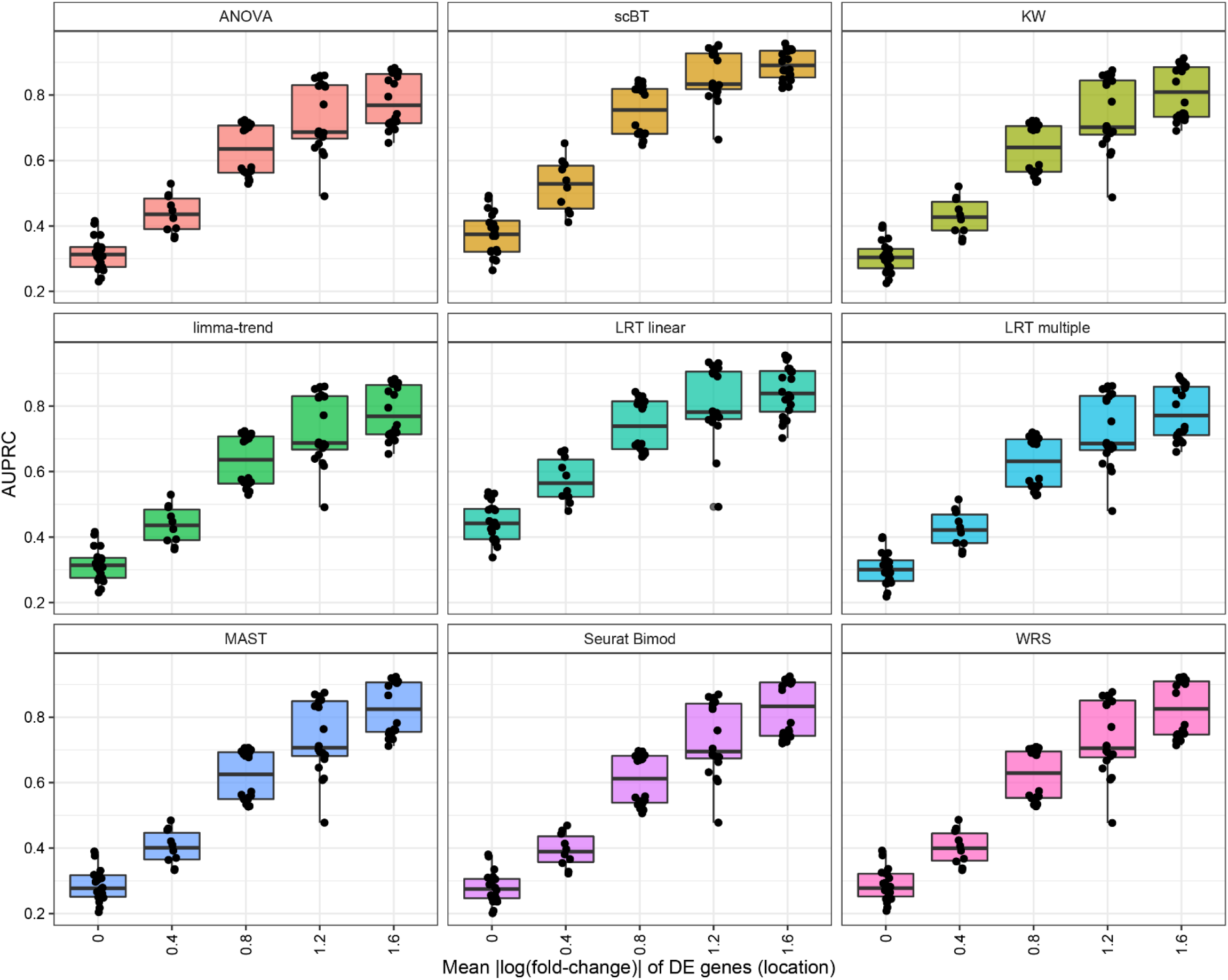
Area under the precision-recall curve (AUPRC) of 9 differential expression test methods for simulated dose response data with varying location of differentially expressed genes. A total of 4,500 cells (500 cells per group) and 5,000 genes were simulated across 9 dose groups with a probability of being differentially expressed of 10%, of which 50% were repressed. Differential expression fold-change scale was 0.4. Box and whisker plots represent median and 25^th^ and 75^th^ percentile, and minimum and maximum values for 10 replicate simulations.

**Supplementary Figure 23:**
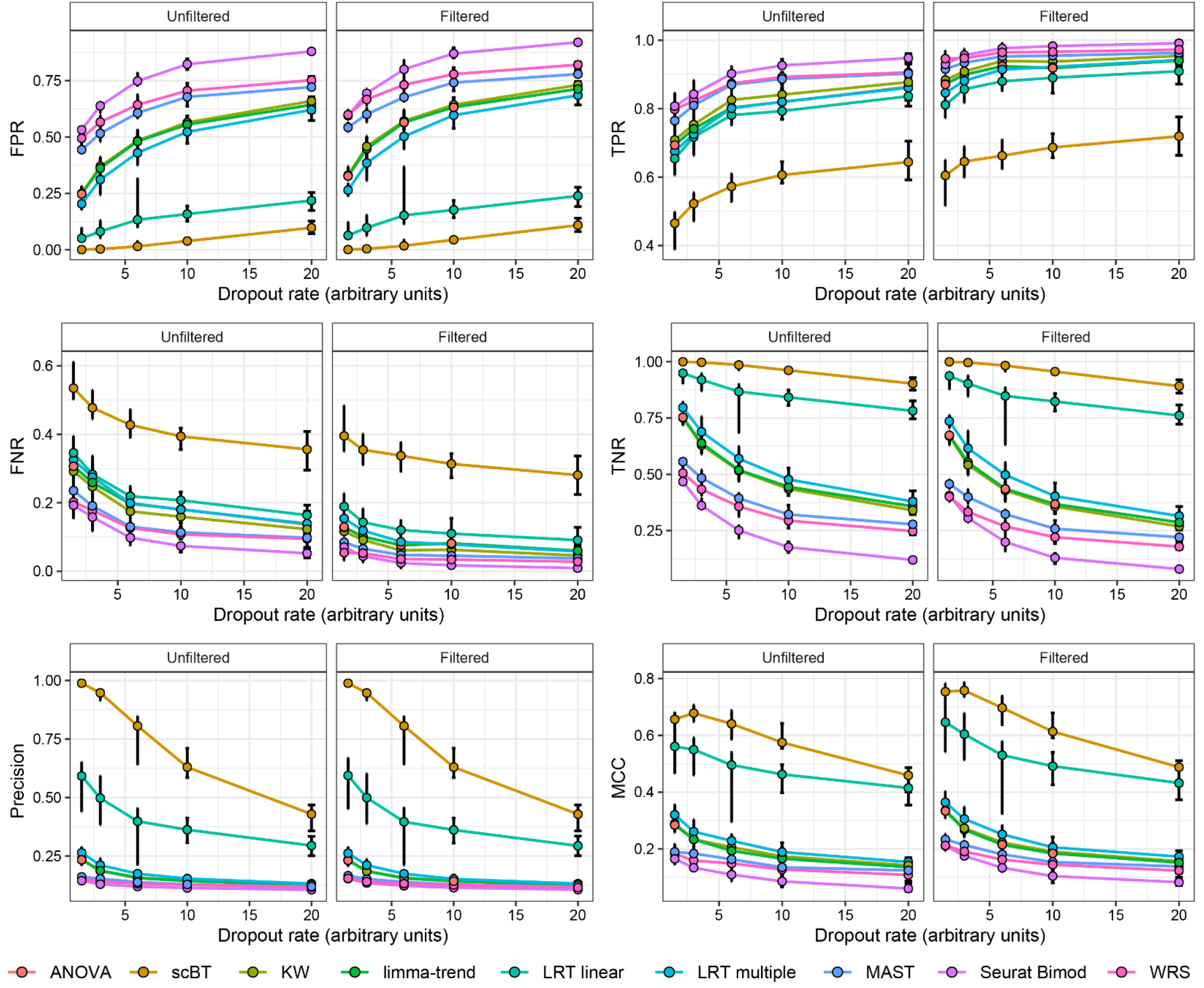
Benchmarking metrics of 9 differential expression test methods for simulated dose response data with varying relationship between mean expression and percent zeroes. A total of 4,500 cells (500 cells per group) and 5,000 genes were simulated across 9 dose groups with a probability of being differentially expressed of 10%, of which 50% were downregulated. Differential expression fold-change location and scale were 0.8 and 0.4, respectively. Given a ground truth from simulation outputs, false positive rates (FPR), true positive rates (TPR), false negative rates (FNR), true negative rates (TNR), precision, and Matthews correlation coefficient (MCC) were calculated. Points represent median ± minimum to maximum values for 10 replicate simulations. Dropout rates are calculated as *Percent zeroes* = *a* ∗ *e*^*b*∗*mean log expression*^ + *t* where parameters are shown in **Table S2**.

**Supplementary Figure 24:**
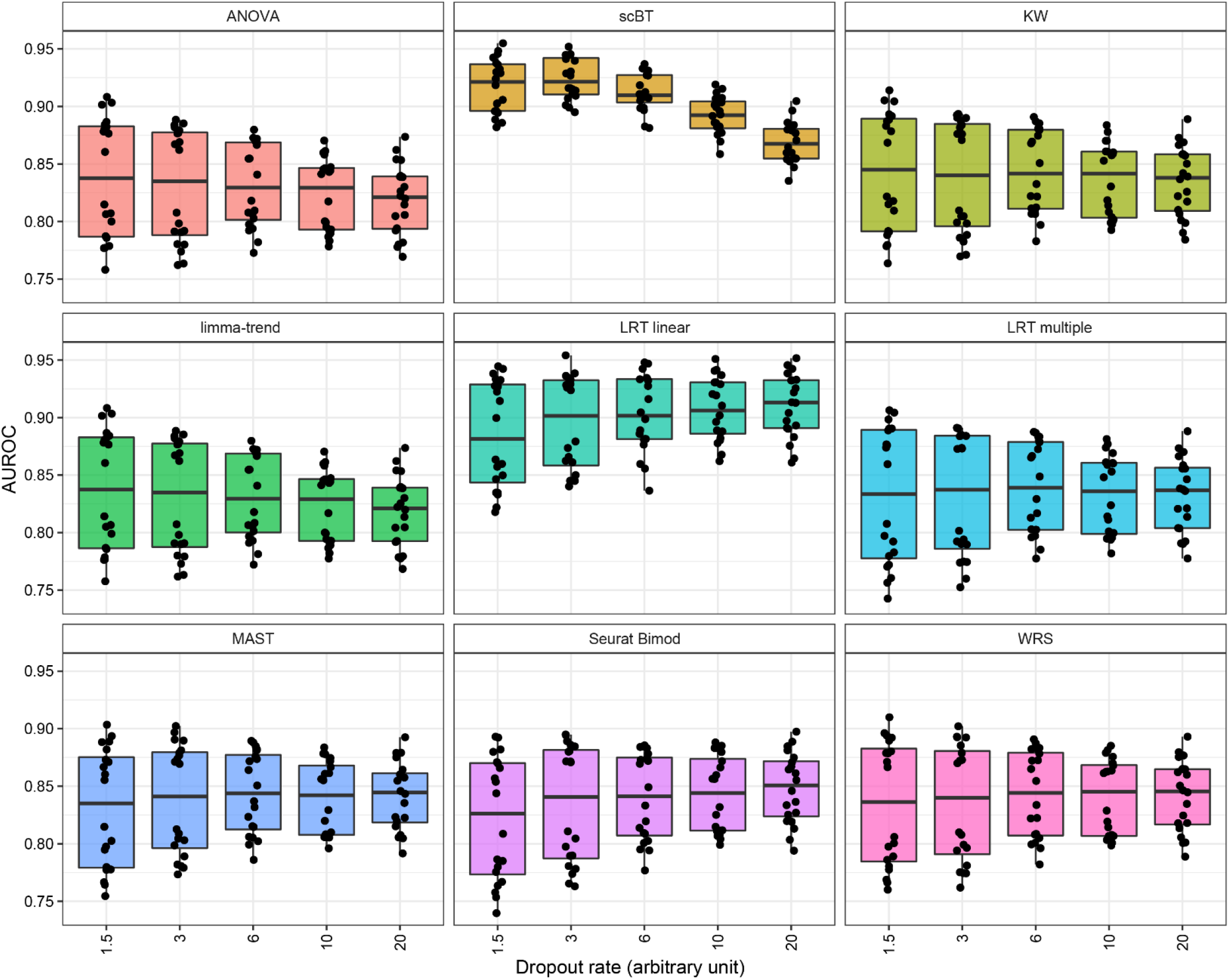
Area under the receiver-operating curve (AUROC) of 9 differential expression test methods for simulated dose response data with varying relationship between mean expression and percent zeroes. A total of 4,500 cells (500 cells per group) and 5,000 genes were simulated across 9 dose groups with a probability of being differentially expressed of 10%, of which 50% were repressed. Differential expression fold-change location and scale were 0.8 and 0.4, respectively. Box and whisker plots represent median and 25^th^ and 75^th^ percentile, and minimum and maximum values for 10 replicate simulations. Dropout rates are calculated as *Percent zeroes* = *a* ∗ *e*^*b*∗*mean log expression*^ + *t* where parameters are shown in **Table S2**.

**Supplementary Figure 25:**
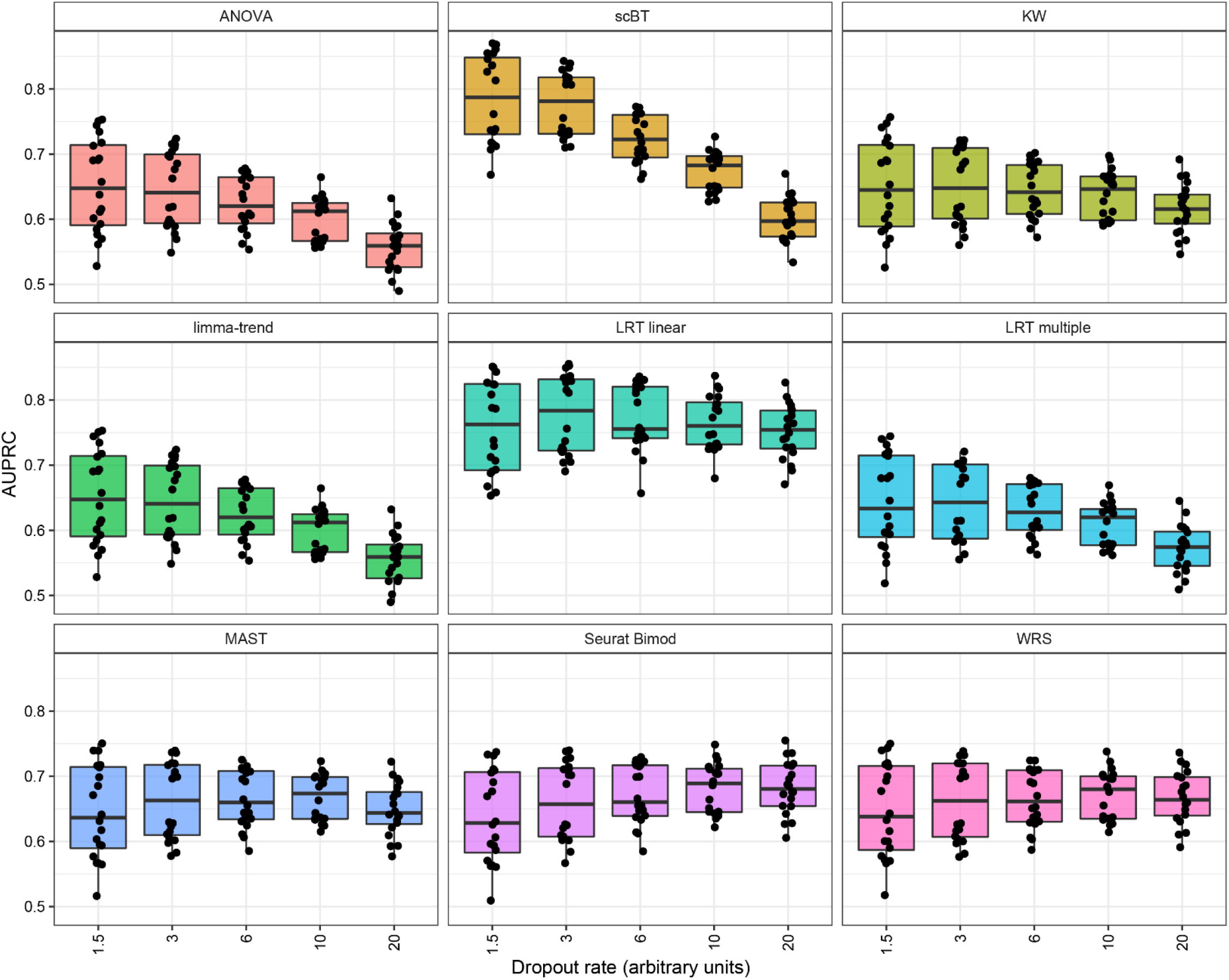
Area under the precision-recall curve (AUPRC) of 9 differential expression test methods for simulated dose response data with varying relationship between mean expression and percent zeroes. A total of 4,500 cells (500 cells per group) and 5,000 genes were simulated across 9 dose groups with a probability of being differentially expressed of 10%, of which 50% were repressed. Differential expression fold-change location and scale were 0.8 and 0.4, respectively. Box and whisker plots represent median and 25^th^ and 75^th^ percentile, and minimum and maximum values for 10 replicate simulations. Dropout rates are calculated as *Percent zeroes* = *a* ∗ *e*^*b*∗*mean log expression*^ + *t* where parameters are shown in **Table S2**.

**Supplementary Figure 26:**
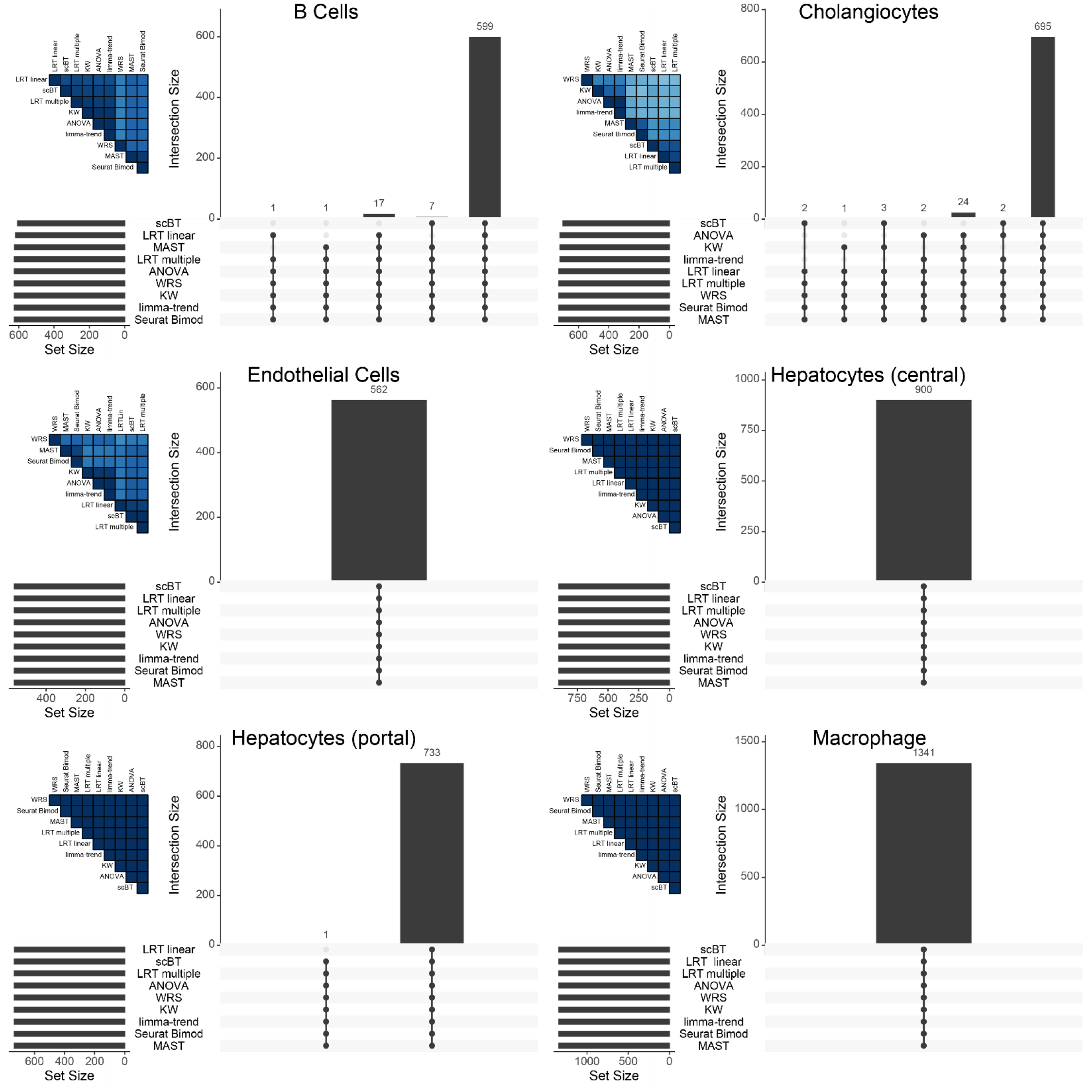
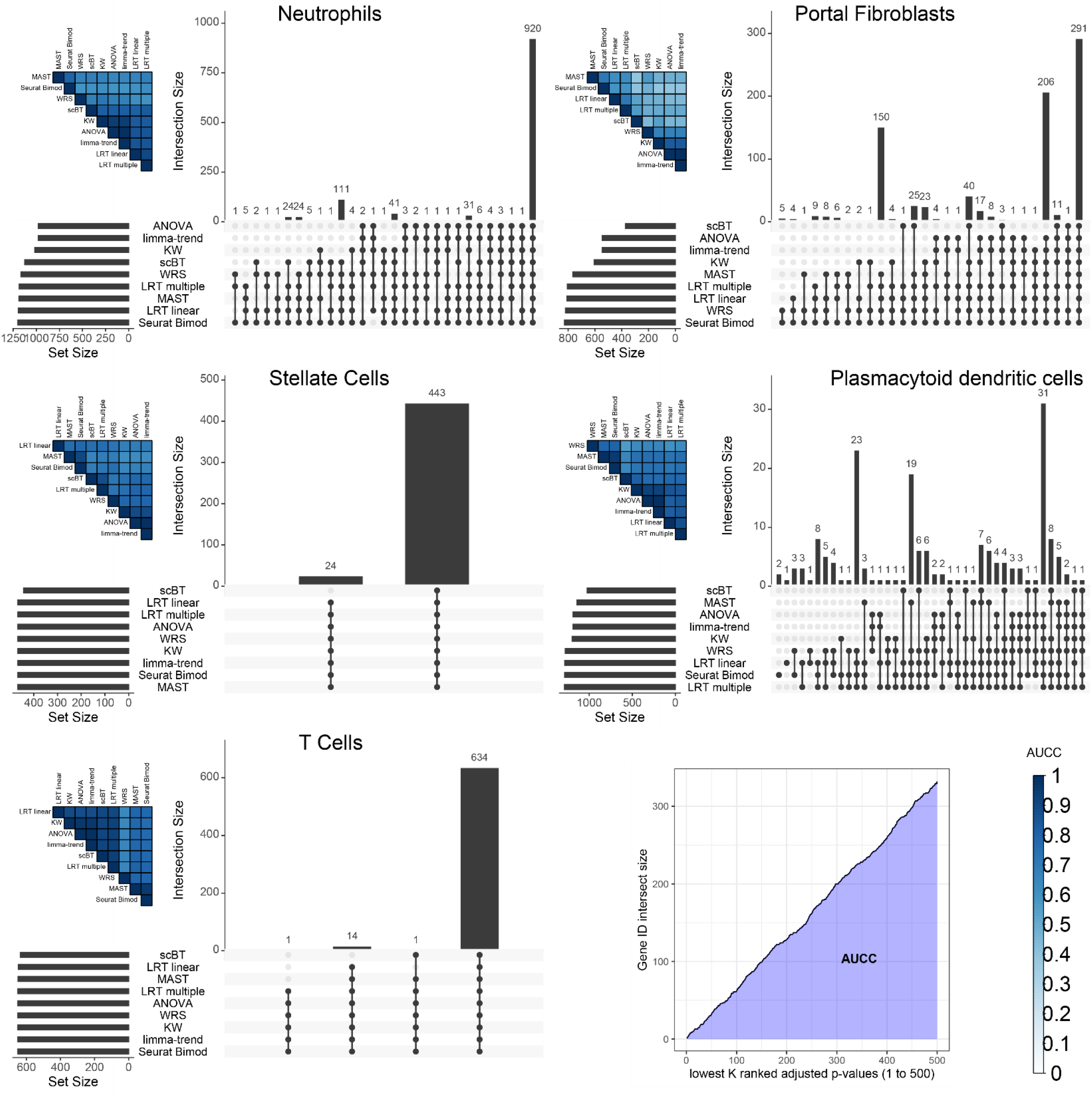
Comparison of differential gene expression analysis of hepatic single-nuclei RNA sequencing data from male mice gavaged with sesame oil vehicle control or 0.01 – 30 µg/kg TCDD every 4 days for 28 days. Each panel represents a distinct cell type showing the intersection of differentially expressed genes (vertical bars) for each combination of tests (filled circles) and total number of differentially expressed genes (horizontal bars). Intersect sizes are displayed on top of vertical bars. The tile plot in the upper left represents the area under the concordance curve (AUCC) calculated in the bottom right panel (page 2). A higher score indicates stronger agreement in the lowest 500 ranked adjusted *p*-values. Genes were considered differentially expressed when expressed in at least 5% of cells in any dose group and with a |fold-change| ≥ 1.5.

**Table S3.**
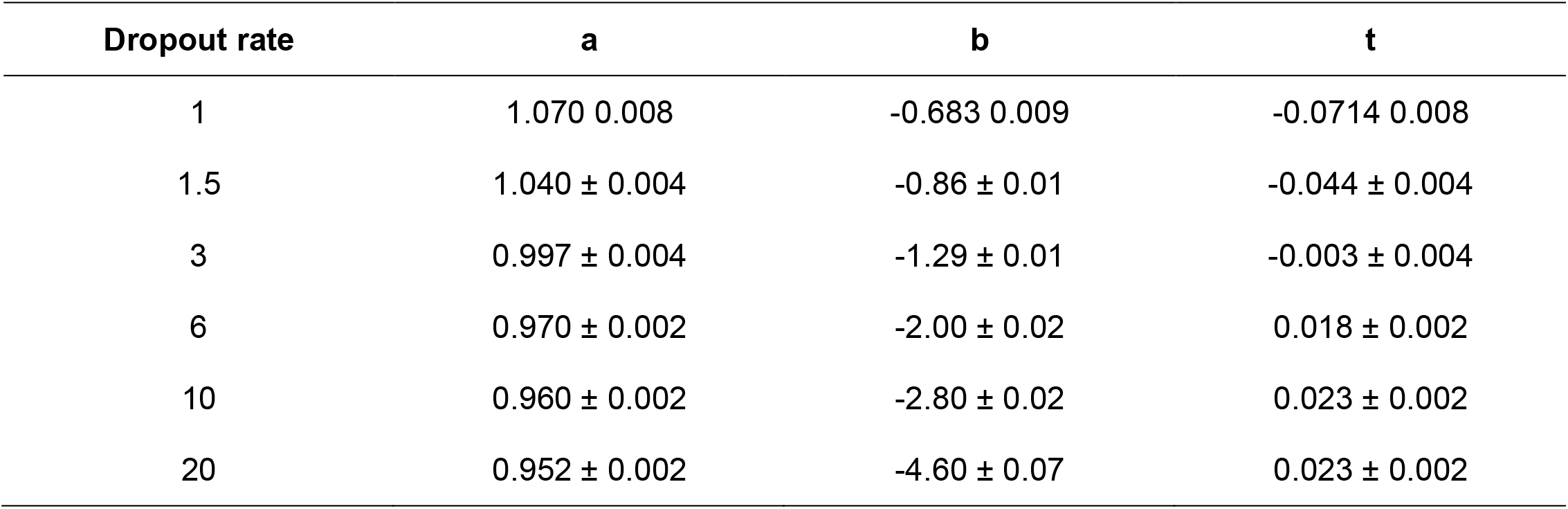
Parameters for modeling of dropout rate calculated using the equation *Percent zeroes* = *a* ∗ *e*^*b*∗*mean log expression*^ + *t*.

| Dropout rate | a | b | t |
| --- | --- | --- | --- |
| 1 | 1.070 0.008 | -0.683 0.009 | -0.0714 0.008 |
| 1.5 | 1.040 ± 0.004 | -0.86 ± 0.01 | -0.044 ± 0.004 |
| 3 | 0.997 ± 0.004 | -1.29 ± 0.01 | -0.003 ± 0.004 |
| 6 | 0.970 ± 0.002 | -2.00 ± 0.02 | 0.018 ± 0.002 |
| 10 | 0.960 ± 0.002 | -2.80 ± 0.02 | 0.023 ± 0.002 |
| 20 | 0.952 ± 0.002 | -4.60 ± 0.07 | 0.023 ± 0.002 |

## Supplementary Material

### 1 Derivation of the marginal likelihoods under the null and alternative hypothesis for scBA

Consider the following K-sample test,

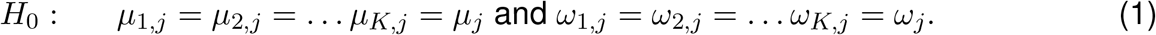

versus the alternative

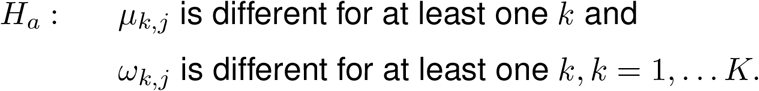

Given this model structure we sssume that a priori, given 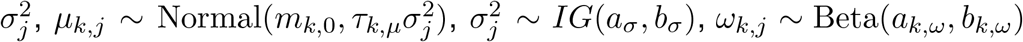, where *IG* is the inverse gamma distribution with shape *a*_*σ*_ and scale *b*_*σ*_ and *m*_*k*,0_, *τ*_*k,µ*_, *a*_*σ*_, *b*_*σ*_, *a*_*k,ω*_, *b*_*k,ω*_ are the hyperparameters. Now, let’s assume that data are collected under K conditions, and denote the data by *D*_*k,o*_ ≡ {(*Y*_*k,i,j*_, *R*_*k,i,j*_), *i* = 1, …, *n*_*k*_}. The underlying populations for the sample data *D*_*k,o*_ for the k=1,2, …, K, are assumed to be identified by the parameters 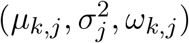. Under the null hypothesis *µ*_1,*j*_ = *µ*_2,*j*_ = … *µ*_*K,j*_ = *µ*_*j*_ and *ω*_1,*j*_ = *ω*_2,*j*_ = … *ω*_*K,j*_ = *ω*_*j*_. We sssume that a priori, given 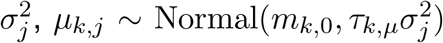, and 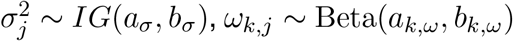, where *m*_*k*,0_, *τ*_*k,µ*_, *a*_*σ*_, *b*_*σ*_, *a*_*k,ω*_, *b*_*k,ω*_ are the hyperparameters. Now we calculate the marginal likelihood under the null hypothesis and alternative hypothesis. Under the null hypothesis the marginal likelihood is

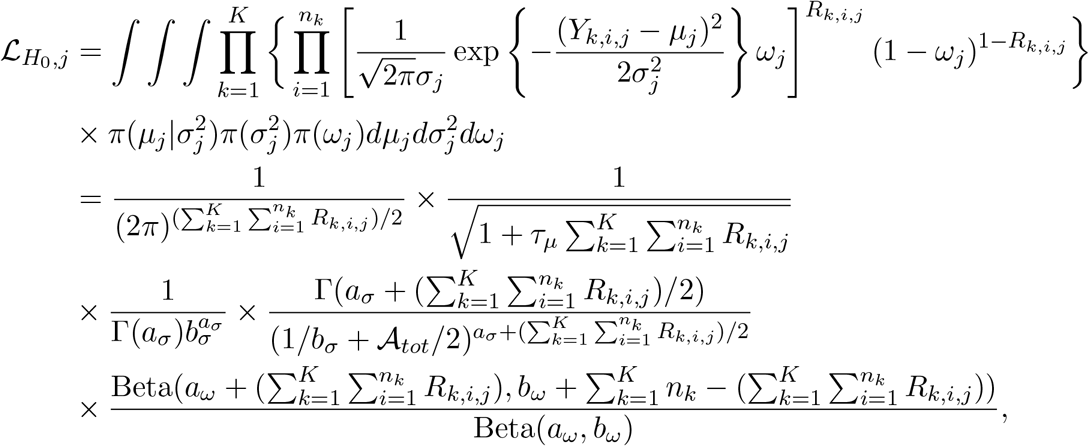

where

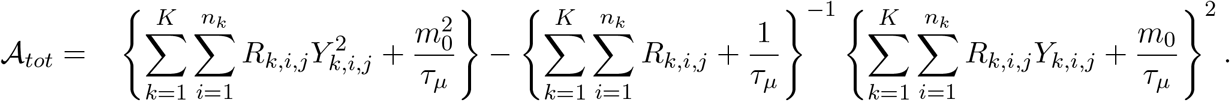

Under the alternative hypothesis we compute the marginal likelihood without any restriction on the K means *µ*_*k,j*_ and the zero inflation parameter *ω*_*k,j*_; *k* = 1, 2, … *K*. Particularly, we assume that 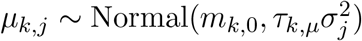, and 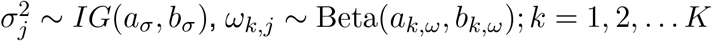. Now,

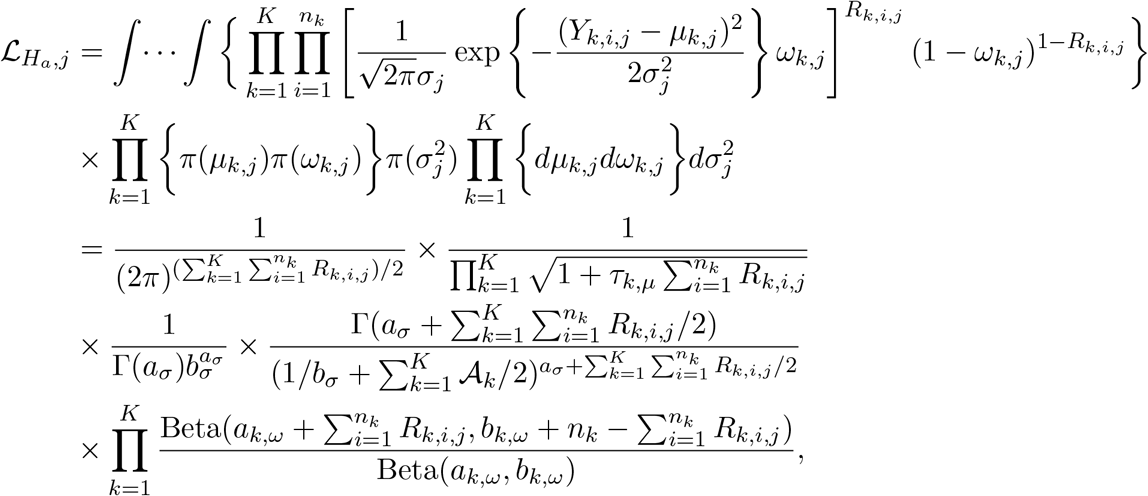

where

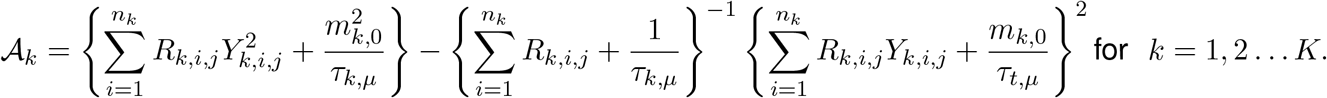

The ratio of the marginal likelihood from *H*_0_ to *H*_*a*_ is

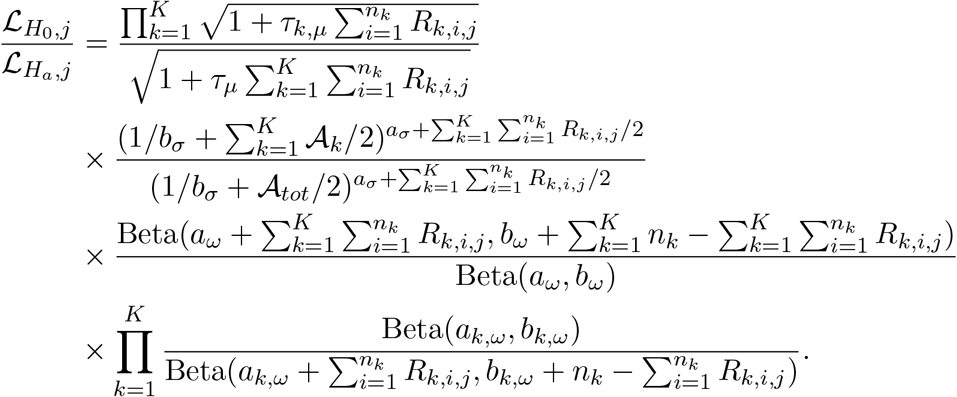

The Bayes factor can be thus be defined as

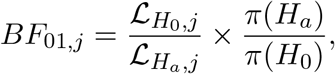

where *π*(*H*_*a*_) and *π*(*H*_0_) are the prior probabilities for the alternative and null model, respectively. To control for multiplicity we adopt the FDR correction approach discussed in^1^. The rejection threshold is estimated in terms of the posterior probabilities of the null hypothesis, *p*(*H*_0,*j*_|*D*_*j*_). For a target FDR *α*, the procedure rejects all hypotheses with *p*(*H*_0,*j*_|*D*_*j*_) *< ζ*, where *p*(*H*_0,*j*_|*D*_*j*_) = (1+ 1*/BF*_01,*j*_)^−1^ and *ζ* is the largest value such that *C*(*ζ*)*/J* (*ζ*) ≤ *α* where, *J* (*ζ*) = {*j* : *p*(*H*_0,*j*_|*D*_*j*_) ≤ *ζ*} and *C*(*ζ*) = ∑_*j*∈*J* (*ζ*)_ *p*(*H*_*o,j*_|*D*_*j*_).

### 2 Derivation of the combined Likelihood Ratio Test Statistic (LRT-multiple)

In this section, we extend the two-sample test proposed by^2^ to a test for *k*-samples. Consider the composite K-sample test

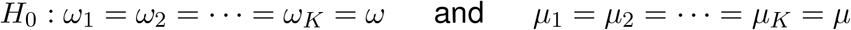

versus the alternative

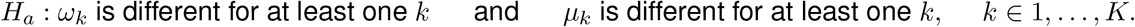

Similar to ANOVA, we assume homogeneity for variance parameter 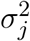. Now, fixing the gene index *j*, the likelihood ratio test can be defined as;

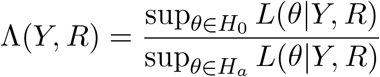

where the likelihood can be written as;

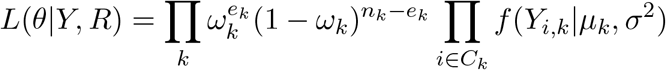

*Y* and *R* represent the gene observation vector and the gene indicator vector across K dose groups and *θ* = {*µ*_*k*_, *σ*^2^, *π*_*k*_, *k* = 1, …, *K*} is the vector of unknown parameters. We define *C*_*k*_ to be the set of cells expressing the gene in group *k* (*i*.*e*.*C*_*k*_ = {*i* : *R*_*ik*_ = 1}) and *e*_*k*_ = ∑_*i*_ *R*_*ik*_ is the cardinality of set *C*_*k*_. Here, *f* denotes the density function of the normal distribution with parameters *µ*_*k*_ and *σ*^2^. Therefore, it follows that the likelihood ratio test can be written as

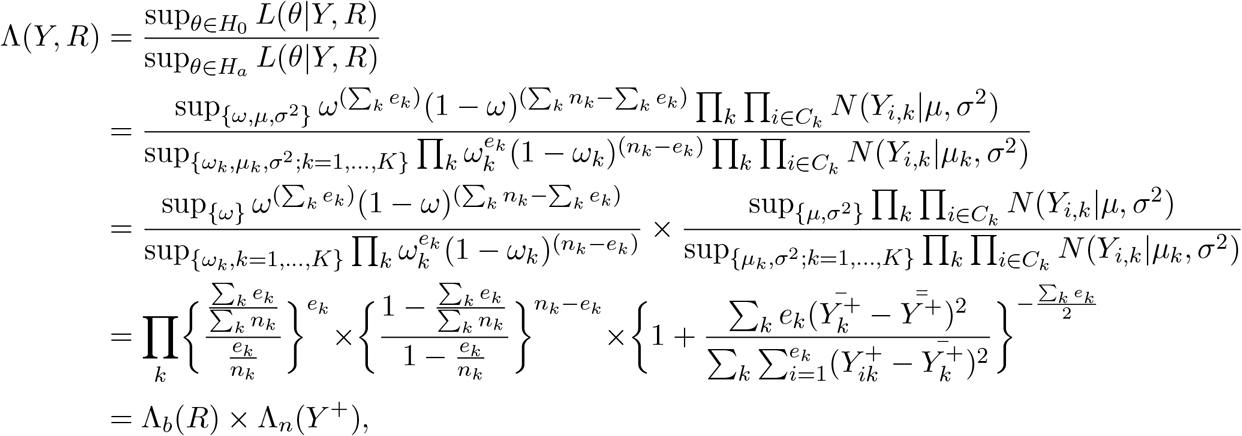

where *N* (·|*µ, σ*^2^) denotes the normal density with mean and variance *µ* and *σ*^2^, Λ_*b*_ is a binomial LRT, Λ_*n*_ is a normal LRT, *Y*^+^ is the set of positive *Y* values, 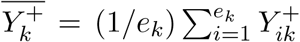 and 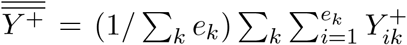. Thus our combined LRT can be computed as the product of a binomial and a normal LRT statistic, both of which can easily be derived using classical statistical theory.

### 3 Derivation of the combined Likelihood Ratio Test Statistic for the linear model setup (LRT-linear)

In this section we extend the combined Likelihood Ratio Test Statistic (LRT-multiple) to a linear model setup. Treating dose (*d*) as a continuous covariate we write *µ*_*ij*_ = *m*_0*j*_ + *d*_*i*_*m*_1*j*_ and logit(*ω*_*ij*_) = *ψ*_0*j*_ +*d*_*i*_*ψ*_1*j*_. Under the null hypothesis the model can be reformulated as *H*_0_ : *µ*_*ij*_ = *m*_0*j*_ and logit(*ω*_*ij*_) = *ψ*_0*j*_. Therefore the likelihood function for gene *j* under the full model can be written as:

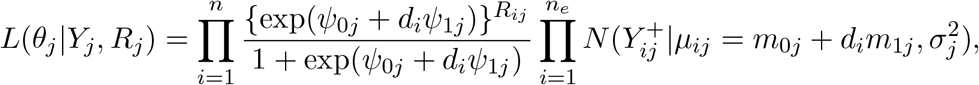

where *R*_*j*_ = *I*(*Y*_*j*_ ≠ 0) denotes the gene expression indicator vector of size 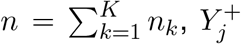 denotes the positively expressed gene observation vector of size *n*_*e*_ = ∑_*k*_ ∑_*i*_ *R*_*ijk*_ and 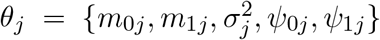 Using the likelihood function described above, the likelihood ratio test can be derived following the same approach detailed in Section 2. Since *R*_*j*_ and *Y*_*j*_ are conditionally independent for each gene *j*, the individuals LRT statistics derived from the logistic and linear regression parts can be summed to obtain an asymptotically *χ*^2^ distribution with the degrees of freedom of the component tests added.

